# Mouse models with human antibody repertoires for inducing multiple lineages of HIV-1 broadly neutralizing antibodies

**DOI:** 10.64898/2026.03.30.715354

**Authors:** Ming Tian, Hwei-Ling Cheng, Jillian Davis, Lily M. Thompson, Aimee Chapdelaine Williams, Marie-Elen Tuchel, Audrey Yin, Lawrence Jianqiao Hu, Xin Lin, Adam Yongxin Ye, Frederick W. Alt

## Abstract

The complementarity determining region (CDR) 1, 2 and 3 of antibodies are the principal antigen contact sites. The heavy chain CDR3 (CDR H3) is highly variable, because it includes random nucleotide additions by the terminal deoxynucleotidyl transferase (TdT). Some broadly neutralizing antibodies (bnAbs) against the human immunodeficiency virus-1 (HIV-1) rely heavily on CDR H3 to recognize conserved epitopes on HIV-1 Envelope (Env) protein. Elicitation of comparable bnAbs is a prime goal of HIV-1 vaccine development, but the shortage of precursor antibodies with suitable CDR H3s in human repertoire is a major challenge for immunogen design. To aid this effort, we generated six mouse models for inducing bnAbs against major HIV-1 Env epitopes. In each mouse model, the immunoglobulin heavy and light chain loci were engineered to predominantly rearrange the germline V, D and J segments of a bnAb lineage. Owing to CDR3 diversity, only a small subset of the V(D)J recombination products may encode variable regions that can engage bnAb epitopes with sufficient affinity to initiate immune response. Therefore, these mouse models can be used to test and optimize immunization strategies to induce bnAbs from rare and diverse precursors in complex antibody repertoires.

## Introduction

Vaccine prevention of the human immunodeficiency virus-1 (HIV-1) infection is hindered by the genetic diversity of circulating viruses. The discovery of bnAbs in individuals living with HIV-1 offered a potential solution to the problem. BnAbs can inhibit a wide spectrum of viral strains by targeting conserved regions of the HIV-1 Env protein. Eliciting bnAbs is a prime goal for HIV-1 vaccine development, and the evolution pathway of bnAbs during infection can serve as a blueprint for vaccine strategies (1). Based on longitudinal studies of bnAb development in human donors, each bnAb lineage originated from an unmutated common ancestor (UCA) that is composed of specific germline immunoglobulin (Ig) V, D, and J segments (2–7). Maturation of the precursor into bnAbs involved extensive somatic hypermutation (SHM) and antigenic selection in the context of co-evolution with the infecting viruses. SHM introduces random mutations into Ig variable regions of antigen-activated B cells in germinal centers, and B cells with improved cognate B cell receptor (BCR) affinities are selected by antigens (8). To recapitulate this process by vaccination, a major limiting factor is the low frequency of bnAb precursors in human antibody repertoire. Many bnAbs rely heavily on CDR H3 for epitope contact (5, 6, 9–19). CDR H3 encompasses V_H_-D and D-J_H_ junctions. During V(D)J recombination, TdT adds random nucleotides to these junctions to form N regions (20). Owing to the stochastic nature of N region synthesis, the frequency of CDR H3s that are suitable for bnAb function is low in the human antibody repertoire (21–24). Furthermore, bnAb epitopes are often shielded by glycans and other structural obstacles (25–27). Reaching these occluded epitopes requires long CDR H3 loops, which are uncommon among human antibodies (23, 28).

Low precursor frequency poses a big challenge for immunogen design. As mentioned above, SHM and antigenic selection of germinal center B cells is a critical part of bnAb evolution. As germline bnAb precursors lack mutations, they generally bind poorly to Env proteins (21, 23, 29–32). To prime bnAb precursor B cells, Envs must be engineered *in vitro* to improve their affinity for germline precursors. After *in vitro* optimizations, immunogens face additional challenges *in vivo*. First, because part of CDR H3 is synthesized in a non-templated manner, the bnAb UCA in the original human donor may not be reproduced in other individuals. Instead, the immunogen would encounter a more diverse range of precursors, with potentially lower affinities than the inferred bnAb precursors that were targeted for *in vitro* optimization. Second, in a complex antibody repertoire, the immunogen will inevitably provoke off-target responses to irrelevant epitopes. When precursor frequency is low, off-target responses can become dominant and preclude bnAb development. To address these concerns, immunogens need to be evaluated and refined by immunization experiments in animal models before human application. To this end, we have generated a panel of mouse models that express diverse human antibodies relevant to several bnAb lineages. These mouse models can be used to test immunogens’ ability to induce bnAbs from infrequent and diverse precursors in complex antibody repertoires.

## Results

### Overview of the humanized IgH and Igκ loci of the mouse models

In these mouse models, we aimed to express bnAb precursor heavy chains (HC) with diverse CDR H3s. Total B cell number in a mouse sets the upper limit of CDR H3 diversity. Because mouse has a much smaller B cell population than human, it is impossible to accommodate the full human antibody repertoire in a mouse. Given this limitation, we engineered the mouse IgH locus to rearrange primarily one set of germline human V_H_, D and J_H_ segments (hV_H_, hD and hJ_H_) of a given bnAb lineage. The strategy can maximize the diversity of CDR H3s of human variable regions that are relevant to the bnAb lineage. To this end, we substituted a hV_H_ for the mouse V_H_5-2 segment and deleted the intergenic control region 1 (IGCRI) on the same IgH allele (Fig 1A). In combination, these modifications can lead to predominant rearrangement of the D-proximal hV_H_ (33, 34). We replaced all four mouse J_H_ segments with a single hJ_H_, which became the only option for V(D)J rearrangements on this IgH allele. We positioned the hD segment immediately upstream of the hJ_H_, in place of the mouse DQ52 segment. A previous study showed that the J_H_-proximal DQ52 was the most frequently utilized D segment in the context of IGCRI deletion (34); we expected this preference would apply to the hD segment at the same position. Furthermore, after hD and hJ_H_ are joined, the recombination product (hD/J_H_) cannot be deleted by secondary D-to-J_H_ rearrangements, as there are no downstream J_H_ segments. This and additional mechanisms can increase the representation of hD/J_H_ in the repertoire (35). Overall, the organization of the humanized IgH allele is analogous to that in the previous VRC01 mouse model (34), except for the use of different hV_H_ and hJ_H_ segments and the addition of hD segment.

**Fig 1.**
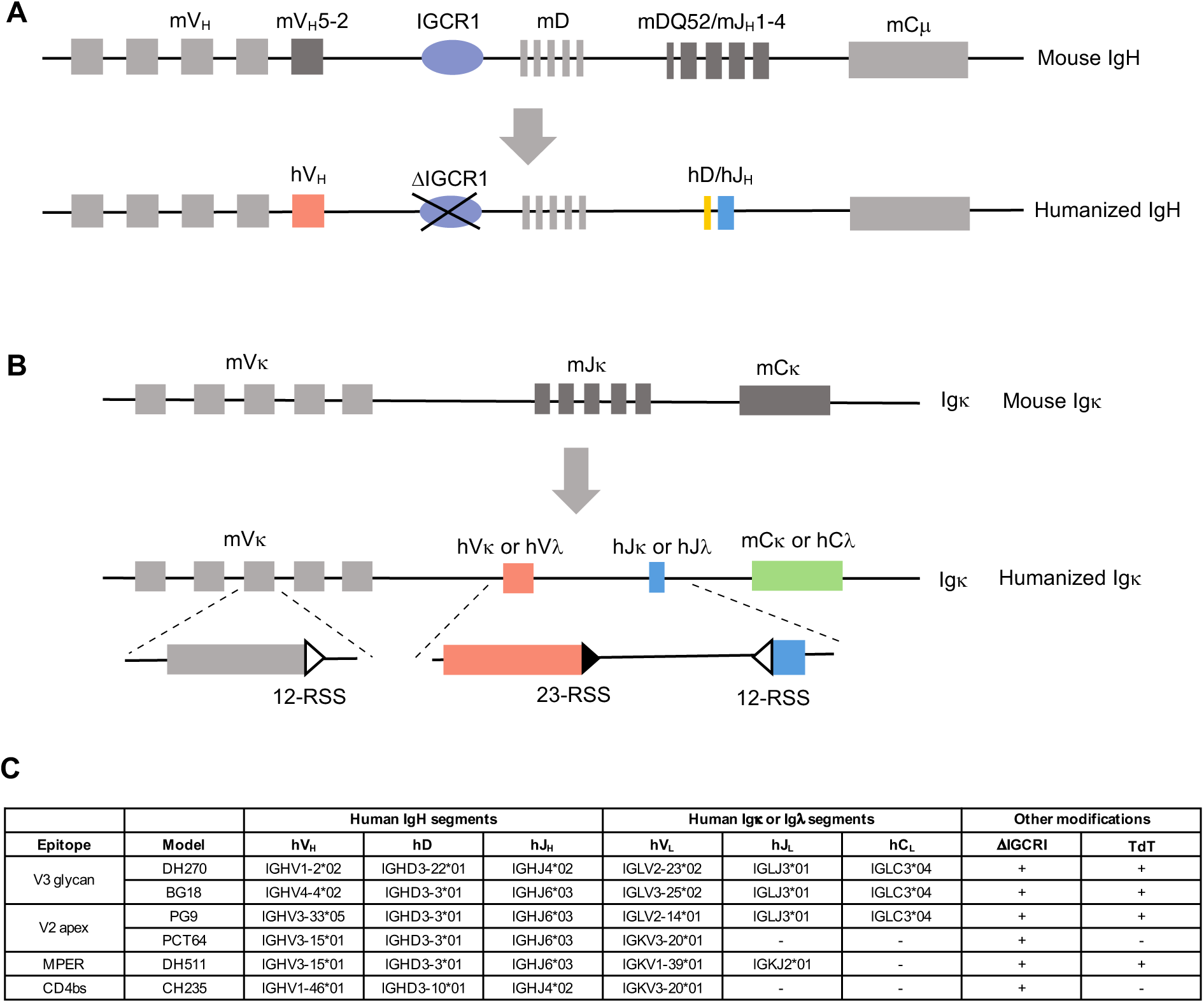
Diagram of the humanized IgH and Igκ allele in the mouse models. (A) Steps in generating the humanized IgH allele. (B) Illustration of the recombination cassette for expressing human LCs in the Igκ allele. (C) Human Ig segments and other genetic modifications in the mouse models. Abbreviations: MPER, membrane proximal external region; CD4bs, CD4 binding site.

To express bnAb precursor light chain (LC), we constructed a recombination cassette that included the germline human V (hV_L_) and J segment (hJ_L_) of a bnAb LC. The cassette was integrated into the mouse Jκ locus, in place of all five mouse Jκ segments (Fig 1B). In the cassette, the hV_L_ segment was flanked by a recombination signal sequence with 23-base pair spacer (23-RSS), whereas the hJ_L_ segment was associated with a 12-RSS. Based on the 12-23 rule of V(D)J recombination (36), the 12-RSS hJ_L_ segment can only recombine with the 23-RSS hV_L_, but not the 12-RSS mouse Vκ segments, which lie upstream of the cassette. Consequently, the humanized Igκ allele should express exclusively human variable regions, despite the presence of intact mVκ cluster. The method, designed to give more dominant expression of the hV_L_ and hJ_L_ in the cassette, differs from the strategy for human LC expression in the VRC01 mouse model, in which a hVκ segment substitutes for a mouse Vκ segment and is expressed in a smaller fraction (up to 10%) of the Igκ light chains (37).

Because Igκ is utilized in most mouse B cells, we incorporated the recombination cassette into the mouse Jκ locus, irrespective of human LC isotype. However, when the human variable region belongs to the Igλ isotype, expression of human Igλ variable region in association with mouse Igκ constant region (mCκ) would generate a hybrid Igλ/κ light chain, which may not fold properly. To address this concern, we replaced mCκ with human Igλ constant region (hCλ3), when the recombination cassette contained human Vλ and Jλ segments. In addition to human Ig segments, a human TdT (hTdT) transgene was incorporated into the *Rosa* locus, as in the VRC01 mouse model (37). Because TdT is expressed during light chain rearrangement in human but not mouse pre-B cells (38, 39), mouse IgL contains less diverse CDR L3s (37). Thus, hTdT transgene was used to allow human-like diversification of CDR L3 in the mouse models (37).

Based on the design described above, we made six mouse models for bnAbs that cover the major sites of vulnerability on the Env protein, including V2 apex (PG9 (40) and PCT64 (12)), V3 glycan, (BG18 (14) and DH270 (5)), Membrane Proximal External Region (MPER) (DH511 (19)), and CD4 binding site (CH235 (3)) (Fig 1C and S1 Table). Some of these bnAbs and their corresponding mouse models share common Ig variable region gene segments. Specifically, BG18 (14), PG9 (40), PCT64 (12) and DH511 (19) antibodies all utilize hD3-3 and hJ_H_6 segments, which together can form long CDR H3s; accordingly, the hD3-3 and hJ_H_6 segments were incorporated into all four mouse models. The BG18, PG9, PCT64 and DH511 mouse models are in fact extensions of an earlier mouse model that we generated and contains just hD3-3 and hJ_H_6 but without any hV_H_ segments (30). In addition to hD3-3 and hJ_H_6, the IGHV3-15*01 (hV_H_3-15) segment is also used by both PCT64 and DH511 antibodies; thus, the IgH alleles of these two mouse models were identical. The IGKV3-20*01 (hVκ3-20) segment in PCT64 LC (12) and the hIGKV3-15*01 (hVκ3-15) segment in CH235 LC (3) have 93.4% sequence identity. A hVκ3-20 rearranging Igκ allele was generated for a VRC01 mouse model (37), and we incorporated this allele into the PCT64 and CH235 models by mouse breeding. The original hVκ3-20 allele also contains a hTdT transgene at the *Rosa* locus (37), and the two loci should normally co-segregate through germline transmission. Unexpectedly, the final PCT64 and CH235 mouse models contained hVκ3-20 but not the hTdT transgene (S1 Fig). The cause is presently unknown; one possibility is that, during germline transmission, meiotic crossover separated the two loci, which are 42x10^6^ bp apart. The loss of hTdT is unlikely to affect the use of these mouse models, as discussed later.

### Characterization of splenic B cells in the mouse models

We used flow cytometry to analyze splenic B cells in the mouse models. For this analysis, all the mouse models were homozygous for the humanized IgH and Igκ alleles, and 129SVE mice represented a wildtype control. We quantified total B (B220^+^) and T cells (CD3^+^) in all the mouse lines (Fig 2A, 2D, S2A and S2B Fig). Since T cells were not modified in the mouse models, we used them as internal reference to compare B cell frequencies in different mouse lines. The spleen of 129SVE mice had about equal proportions of B and T cells, corresponding to a B/T cell ratio of about 1. Relative to 129SVE mouse, the B/T cell ratios of the mouse models were lower, reflecting reduction in B cell frequency: DH270 (5) (73%), PG9 (68%), PCT64 (51%), CH235 (44%), DH511 (37%), BG18 (29%) (Fig 2D); correspondingly, splenic B cell numbers were reduced to similar degrees (S2C Fig). The severity of the phenotype varied for different mouse models, likely associated with the propensity of the human Ig variable region gene segments in each mouse model to form autoreactive antibodies or unstable HC/LC pairs, which are subject to negative selection during B cell development.

**Fig 2.**
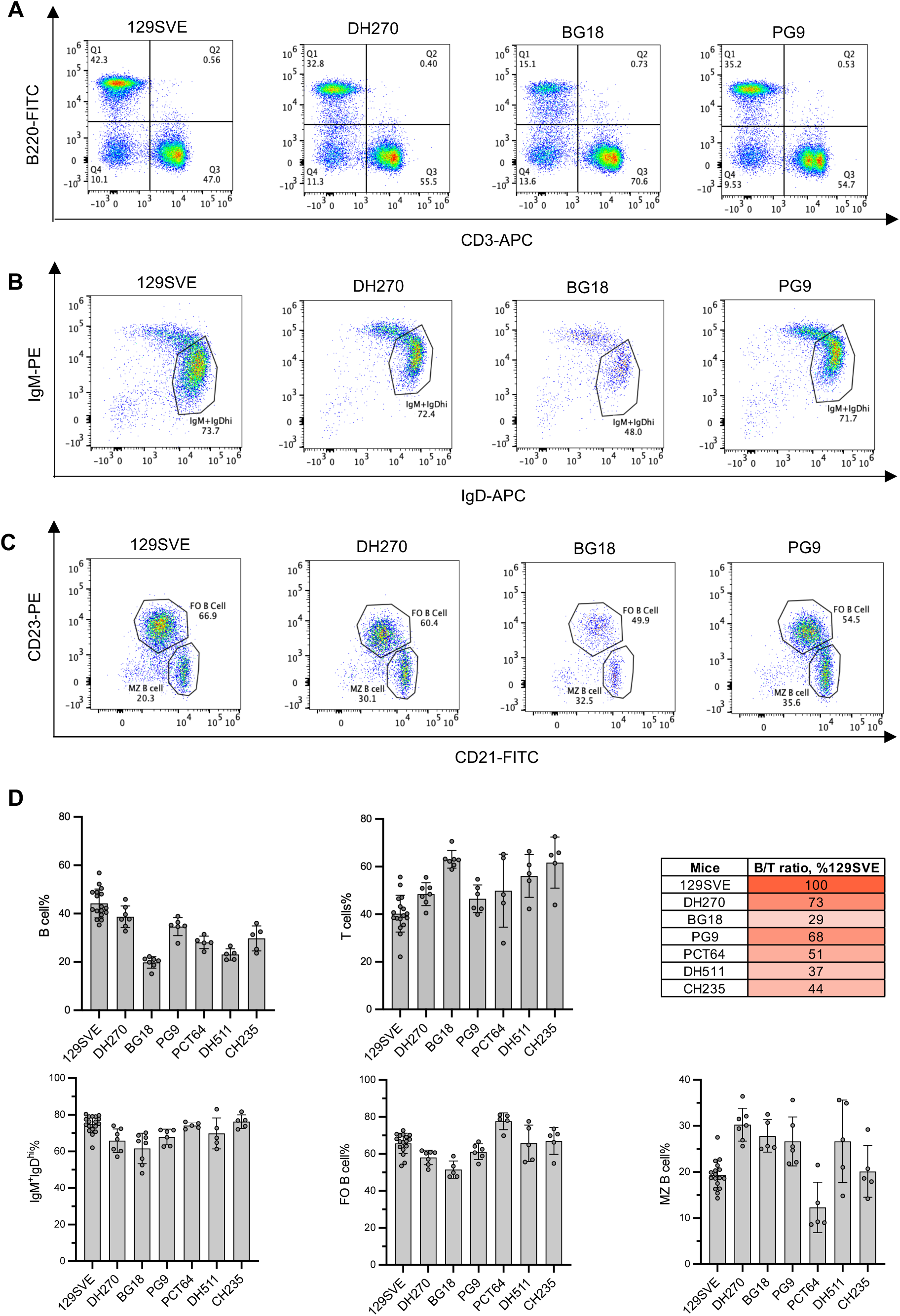
Flow cytometric analysis of splenic B cells of the mouse models. FACS plots for 129SVE mouse, and DH270, BG18, PG9 models are shown in this figure. FACS plots for PCT64, DH511 and CH235 models, including gating schemes, are in S2 and S3 Fig. (A) Total B cells and T cells. Gating scheme: lymphocyte>live cells>single cells>B220/CD3 (S2A Fig). Q1: B cells (B220^+^CD3^-^); Q3: T cells (B220^-^CD3^+^). (B) Surface IgM and IgD expression on B cells. Although some BCRs contain human variable regions, all express mouse μ or δ constant regions, which are the targets of the staining antibodies. Gating scheme: lymphocyte>live cells>single cells>B220^hi^ B cells>IgM/IgD (S2A and S2D Fig). The IgM^+^IgD^hi^ gate contains mature follicular B cells. (C) Follicular and marginal zone B cells. Gating scheme: lymphocyte>live cells>single cells> B220^hi^CD93^lo^ mature B cells>CD21/CD23 (S2A and S3C Fig). Follicular (FO) B cells are B220^hi^CD93^lo^CD21^lo^CD23^hi^; marginal zone (MZ) B cells are B220^hi^CD93^lo^CD21^hi^CD23^lo^. (D) Summary of FACS analysis. The bar graphs and table are based on the FACS analysis shown in panels A-C of this figure, S2 and S3 Fig. Each dot in the graph represents one mouse. The height of the bar corresponds to the mean. Error bar represents standard deviation. Based on this data, the average B/T ratio for each mouse model was calculated. The average B/T ratio for each mouse model was divided by that of the 129SVE mouse, and resulting value is listed in the table to represent the B cell frequencies of each mouse model relative to 129SVE mouse (B/T ratio, %129SVE).

Nascent B cells from bone marrow complete maturation from transitional B cells to follicular B cells in the spleen. The maturation process correlates with a gradual drop in surface IgM level and reciprocal elevation of IgD expression (41). B cells from the mouse models exhibited largely normal IgM and IgD profiles. The proportions of IgM^+^IgD^hi^ population, which contains mature follicular B cells, were comparable for all the mouse lines (Fig 2B, 2D, and S3A Fig). None of the splenic B cell samples showed obvious signs of maturation block, such as accumulation of IgM^hi^IgD^lo^ transitional B cells (41) or IgM^lo^IgD^lo^ anergic B cells (42). Within the mature B cell population (B220^hi^CD93^lo^) (43) (S3B Fig), the distribution of follicular B cell (CD21^lo^CD23^hi^) and marginal zone B cell (CD21^hi^CD23^lo^) (44) was comparable for all mouse lines (Fig 2C, 2D, S3C Fig).

The B220^+^ population (Fig 2A and S2B Fig) included total B cells. Normally, most splenic B cells express uniformly high levels of B220, as seen in 129SVE mouse, DH270, PG9, PCT64, DH511, and CH235 models (Fig 2A, S2B Fig), but B cells in the BG18 model deviated from the norm, as a substantial fraction of B cells exhibited lower levels of B220 staining (Fig 2A). The B220 low (B220^lo^) fraction was more distinct on B220/FSC plot, owing to its slightly larger size than the B220-high (B220^hi^) population (S2D Fig); about 33% of B cells in the BG18 model fell into this category (S2D Fig). This B220^lo^ population expressed typical B1 B cell markers: CD5^+^, CD43^+^ (S2E and S2F Fig) and IgM^+^IgD^lo^ (S3D Fig) (45, 46). B1 B cells represent a distinct lineage from conventional B2 B cells, such as the predominant follicular B cells in spleen (47). B1 B cells arise during fetal hematopoiesis and reside predominantly in the peritoneal cavity of adult mice (48). The presence of such a sizable B1 B cell population in the spleen of the BG18 model was unusual. B1 B cell fate is influenced by BCR specificity. The BCRs of B1 B cells tend to be polyreactive, and interaction with self-antigen may play a role in directing B cell differentiation toward the B1 lineage (49). It remains to be determined whether, and what, polyreactive BCRs underlie the expansion of B1 B cells in the BG18 model.

B1 and marginal zone B cells have restricted BCR repertoires and primarily perform innate-like immune functions. By contrast, follicular B cells express diverse BCRs, respond to a wide range of antigens, carry out affinity maturation in germinal centers and maintain immune memory. Thus, follicular B cells are the most relevant B cell subset for bnAb induction. Because the mouse models showed total B cell reduction, they should also have proportionally smaller follicular B cell compartments than wild-type mice. As shown below, human variable regions dominate the peripheral B cell repertoire, and the human BCR repertoire in each mouse model is focused on one bnAb lineage. Thus, despite the reduction in B cell compartment, each mouse model should still have ample B cells that express BCRs relevant to a type of bnAb.

### IgH repertoire analysis of the mouse models

We performed HTGTS-rep-seq (50) to analyze V(D)J rearrangements in splenic B cells of the mouse models. Because this technique quantifies V(D)J rearrangements on genomic DNA, its output is not influenced by Ig transcript abundance. We used a primer downstream of the hJ_H_ segment to initiate library synthesis for Illumina sequencing. Since the humanized IgH allele contained only one hJ_H_ segment, the libraries included all V(D)J rearrangements on this allele. We focused on productive rearrangements, in which the V, D, J segments were joined in-frame and uninterrupted by stop codons. Due to allelic exclusion, each mature B cell should contain only one productive IgH rearrangement.

This analysis showed that the majority of IgH rearrangements involved the hV_H_ segment in each mouse model: hV_H_1-2, 89.4% in the DH270 model; hV_H_4-4, 65.2% in the BG18 model, hV_H_3-33, 84.9% in the PG9 model, hV_H_3-15, 44.4% in the PCT64 model, hV_H_3-15, 58.1% in the DH511 model, and hV_H_1-46, 87.8% in the CH235 model (Fig 3A, 3B, S4A-S4E Fig). The hV_H_ segment was joined mainly to the hD segment: hD3-22, 82.3% in the DH270 model; hD3-3, 87.0% in the BG18 model; hD3-3, 85.9% in the PG9 model; hD3-3, 69.9% in the PCT64 model; hD3-3, 70.8% in the DH511 model; hD3-10, 90.6% in the CH235 model (Fig 3C, 3D and S5A-S5E Fig). The three reading frames (RFs) of the hD segment were represented unevenly in the productive rearrangements (Fig 3E and S5F Fig). The bias likely resulted from antigenic and structural selection in the context of BCR. Because the RF1 of hD3-22 is interrupted by 3 stop codons in the middle, it was not used in productive rearrangements in the DH270 model. The RF1 of hD3-10 has a stop codon at the end, which can be trimmed off during V(D)J recombination. For this reason, RF1 of hD3-10 was still represented in productive rearrangements in the CH235 model. Because of frequent hV_H_/hD joining and the obligate use of hJ_H_, a large fraction of V(D)J rearrangements were composed entirely of human Ig segments: hV_H_1-2/D3-22/J_H_4 (74.0%) in the DH270 model; hV_H_4-4/D3-3/J_H_6 (56.9%) in the BG18 model; hV_H_3-33/D3-3/J_H_6 (73.0%) in the PG9 model; hV_H_3-15/D3-3/J_H_6 (30.9%) in the PCT64 model; hV_H_3-15/D3-3/J_H_6 (41.2%) in the DH511 model; hV_H_1-46/D3-10/J_H_4 (79.6%) in the CH235 model (Fig 3F). Altogether, these results confirmed that the humanized IgH allele in each mouse model rearranges predominantly one set of human V_H_, D and J_H_ segments of a bnAb.

**Fig 3.**
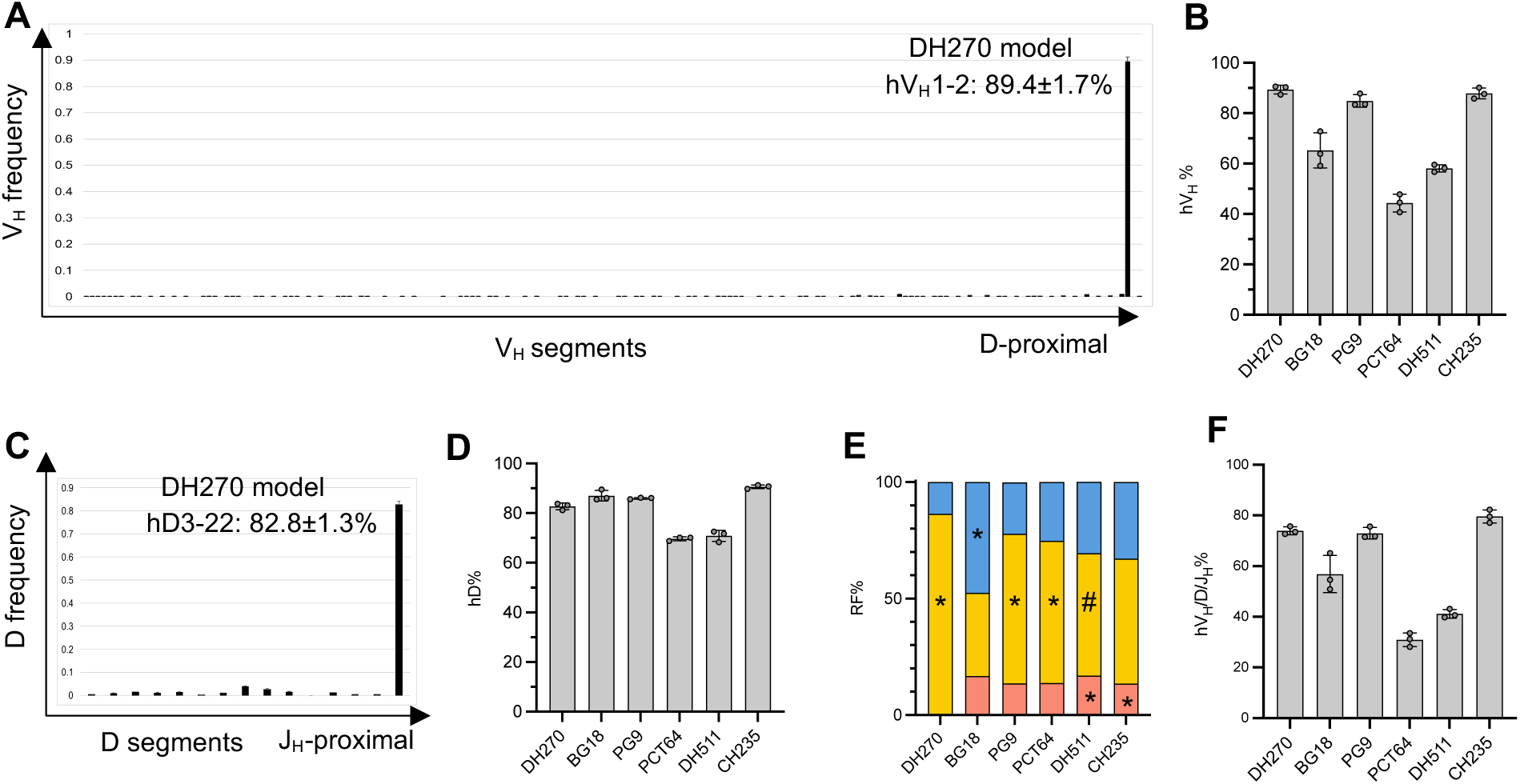
IgH repertoire analysis of splenic B cells from the mouse models. (A) V_H_ usage in the DH270 mouse model. The plot shows V_H_ usage among productive rearrangements. On the x-axis, the V_H_ segments on the humanized IgH allele are arrayed from distal (left) to proximal (right) relative to the D segments. The hV_H_1-2 segment and its usage are shown. The height of the bar corresponds to the mean of three mice. Error bar represents standard deviation. The mouse V_H_ segments are not labeled, because of their large number and minimal usage. The y-axis shows V_H_ usage frequency, which is calculated by dividing the reads of productive rearrangements for each V_H_ by the total reads of productive rearrangements. The plots for the BG18, PG9, DH511, PCT64, CH235 models are in S4 Fig. (B) Summary of hV_H_ usage in the mouse models. The bar graph is based on the data in panel A of this figure and S4A-S4E Fig. The height of the bar equals the mean. The error bar represents standard deviation. (C) D usage of the DH270 model. The plot shows D usage frequency among productive rearrangements that contain hV_H_1-2 and hJ_H_4. The plot is labeled as in panel A. The D usage plots for the BG18, PG9, DH511, PCT64, CH235 models are in S5 Fig. (D) Summary of hD usage in the mouse models. The bar graph is based on the data in panel C of this figure and S5 Fig. (E) D reading frame (RF) usage in productive rearrangements of hV_H_/D/J_H_. Reading frame 1 is in red, reading frame 2 is in yellow, and reading frame 3 is in blue. The reading frame that is used in the bnAb for each mouse model is marked with *. The 10E8 D reading frame is marked with # for the DH511 mouse model. (F) The frequency of hV_H_/D/J_H_ in total productive rearrangements. The frequency is calculated by hV_H_% x hD% x hJ_H_%, based on hV_H_% in panel B, hD% in panel D and hJ_H_%=100%.

Long CDR H3s are necessary for the function of many bnAbs, including DH270, BG18, PG9, PCT64 and DH511. We analyzed the CDR H3 length of the productive human IgH rearrangements in the mouse models. CDR H3 length correlates with D and J_H_ segments. The D segments in these mouse models, hD3-10, hD3-22 and hD3-3, are equal in length; but hJ_H_6 is longer than hJ_H_4. Correspondingly, the IgH rearrangements of hJ_H_6-based mouse models (BG18, PG9, PCT64 and DH511) had longer CDR H3s than those from hJ_H_4-based mouse models (DH270 and CH235) (compare Fig 4A and 4B). We further compared CDR H3 length between mouse model and human. The published human antibody repertoire data were generated with RNA-based methods (23, 28), whereas the IgH repertoire analysis of the mouse models was performed with DNA-based technique (50). For a matching comparison, we used HTGTS-rep-seq to analyze human IgH rearrangements on genomic DNA from peripheral blood B cells (S6 Fig), and we also analyzed C57BL/6 mouse IgH repertoire in parallel (S6 Fig). Since mouse J_H_2 and J_H_3 are equal in length to human J_H_4, we combined mouse J_H_2 and J_H_3-based CDR H3s to compare with human J_H_4-based CDR H3s. Similarly, the CDR H3 distributions of the DH270 and CH235 models were combined into one data set (DH270+CH235). For J_H_6-based CDR H3s, we combined the CDR H3 data from BG18, PG9, PCT64 and DH511 mouse models. Mouse does not have any J_H_ that is as long as human J_H_6. Instead, we used mouse J_H_1 and J_H_4, which are longer than J_H_2 and J_H_3. By superimposing the CDR H3 profiles of human, mouse model, and C57BL/6 mouse, it was apparent that the mouse models had similar CDR H3 length distribution as human, and C57BL/6 mouse had shorter CDR H3s than both (Fig 4C and 4D).

**Fig 4.**
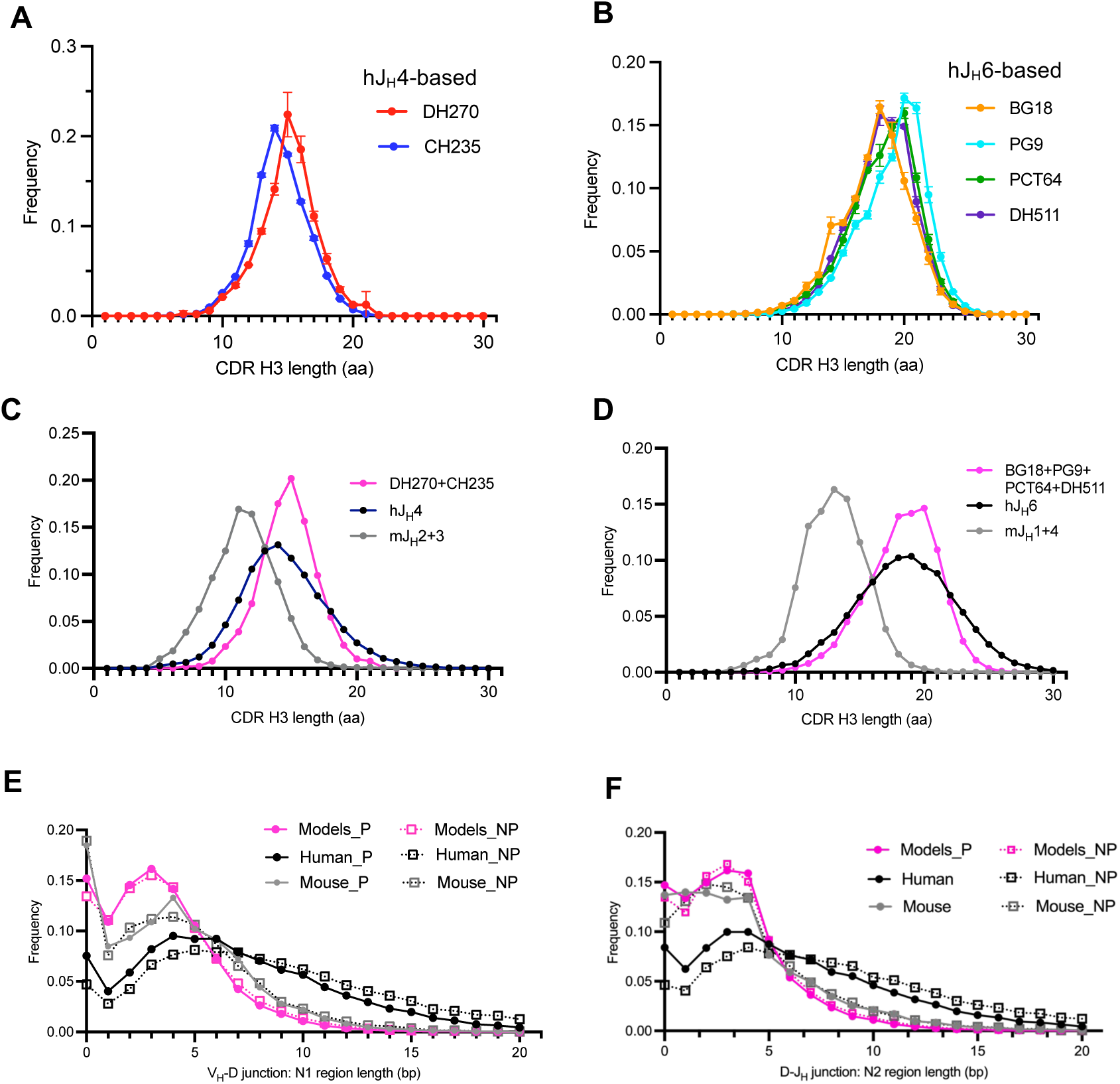
CDR H3 analysis of the mouse models. (A-B) CDR H3 length distribution of productive rearrangements of hV_H_1-2/D3-22/J_H_4 (DH270 model), hV_H_1-46/D3-10/J_H_4 (CH235 model), hV_H_4-4/D3-3/J_H_6 (BG18 model), hV_H_3-33/D3-3/J_H_6 (PG9 model), hV_H_3-15/D3-3/J_H_6 (PCT64 model and DH511 model). The plot is based on the HTGTS-rep-seq analysis in Fig 3, S4 and S5 Fig. The plot for each mouse model is based on analysis of 3 mice, and error bar represents standard deviation. (C-D) CDR H3 length comparison. DH270+CH235 is the average of the two mouse models, based on the data in panel A. BG18+PG9+PCT64+DH511 is the average of the four mouse models, based on the data in panel B. hJ_H_4 and hJ_H_6 are hJ_H_4-based and hJ_H_6-based productive rearrangements in human B cells, based on S6A and S6B Fig. mJ_H_2+3 is the average of mJ_H_2 and mJ_H_3-based productive rearrangements in C57BL/6 mouse B cells, based on S6C and S6D Fig; mJ_H_1+4 is the average of mJ_H_1 and mJ_H_4-based productive rearrangements in C57BL/6 mouse B cells, based on S6E and S6F Fig. (E-F) N1 and N2 region comparison. Models: average of all six mouse models; Human: average of human J_H_4 and J_H_6-based rearrangements; Mouse: average of C57BL/6 mouse J_H_1-J_H_4. P: productive rearrangements; NP: non-productive rearrangements.

Despite overall similarity, the CDR H3 profiles of mouse models and humans showed noticeable differences. Relative to mouse models, human had more CDR H3s beyond 18aa for hJ_H_4-based CDR H3s (Fig 4C) and beyond 22aa for hJ_H_6-based CDR H3s (Fig 4D). Since the CDR H3s of the mouse models already utilized the longest human D and J_H_ segments, the difference may lie in N regions. CDR H3 contains two N regions: N1 between V_H_ and D; N2 between D and J_H_. Consistent with prediction, both N1 and N2 regions in human CDR H3s extended into longer ranges than the counterparts in mouse models and C57BL/6 mouse (Fig 4E and 4F). The difference could reflect intrinsic properties of the V(D)J recombination machinery or functional selection. To differentiate between the two possibilities, we analyzed N regions in non-productive rearrangements, which should not be subject to functional selection. We found the same difference in N region length among non-productive rearrangements (Fig 4E and 4F). These results suggested that the human V(D)J recombination machinery can generate longer N regions than the mouse system. Similar observations have been made in CDR H3 analysis of the Kymouse, which incorporated the complete set of human V_H_, D and J_H_ segments (51, 52). It is unclear why N regions are shorter in mouse than in human. TdT is expressed during IgH rearrangements in mouse pro-B cells and is responsible for N region synthesis in CDR H3 (53, 54). Additionally, some of the rearranging mouse models harbored a hTdT transgene. In the previous VRC01 mouse model, the hTdT transgene had no detectable impact on CDR H3 length (37). Thus, TdT activity may not be the limiting factor for N region length in CDR H3.

### Identification of bnAb signature in the IgH repertoires of the mouse models

Within the human IgH rearrangements in each mouse model, we searched for “bnAb-like” HCs, which were defined as sharing the following bnAb signature: 1) same V_H_, D and J_H_ segment; 2) same CDR H3 length; 3) use of D segment in the same reading frame; 4) start of D segment motif at the same position in CDR H3. The definition is analogous to those in previous bnAb precursor analysis (21–23). Based on these criteria, we found bnAb-like HCs at frequencies of 1.5 x10^-4^ (1 in 6667 B cells) in the DH270 model, 5.1x10^-4^ (1 in 1961 B cells) in the BG18 model, and 2.1x10^-4^ (1 in 4762 B cells) in the PCT64 model (Fig 5A-C and 5G). In these bnAb-like CDR H3s, the N1 and N2 regions, which flank the D segment, were diverse. Unlike DH270, BG18 and PCT64, the CDR H3 of CH235UCA is relatively short (15aa) and contains only 2 residues from the hD3-10 segment (4). Consequently, the frequency of CH235-like CDR H3 was high: 4.2x10^-2^ (1 in 24 B cells) (Fig 5D and 5G).

**Fig 5.**
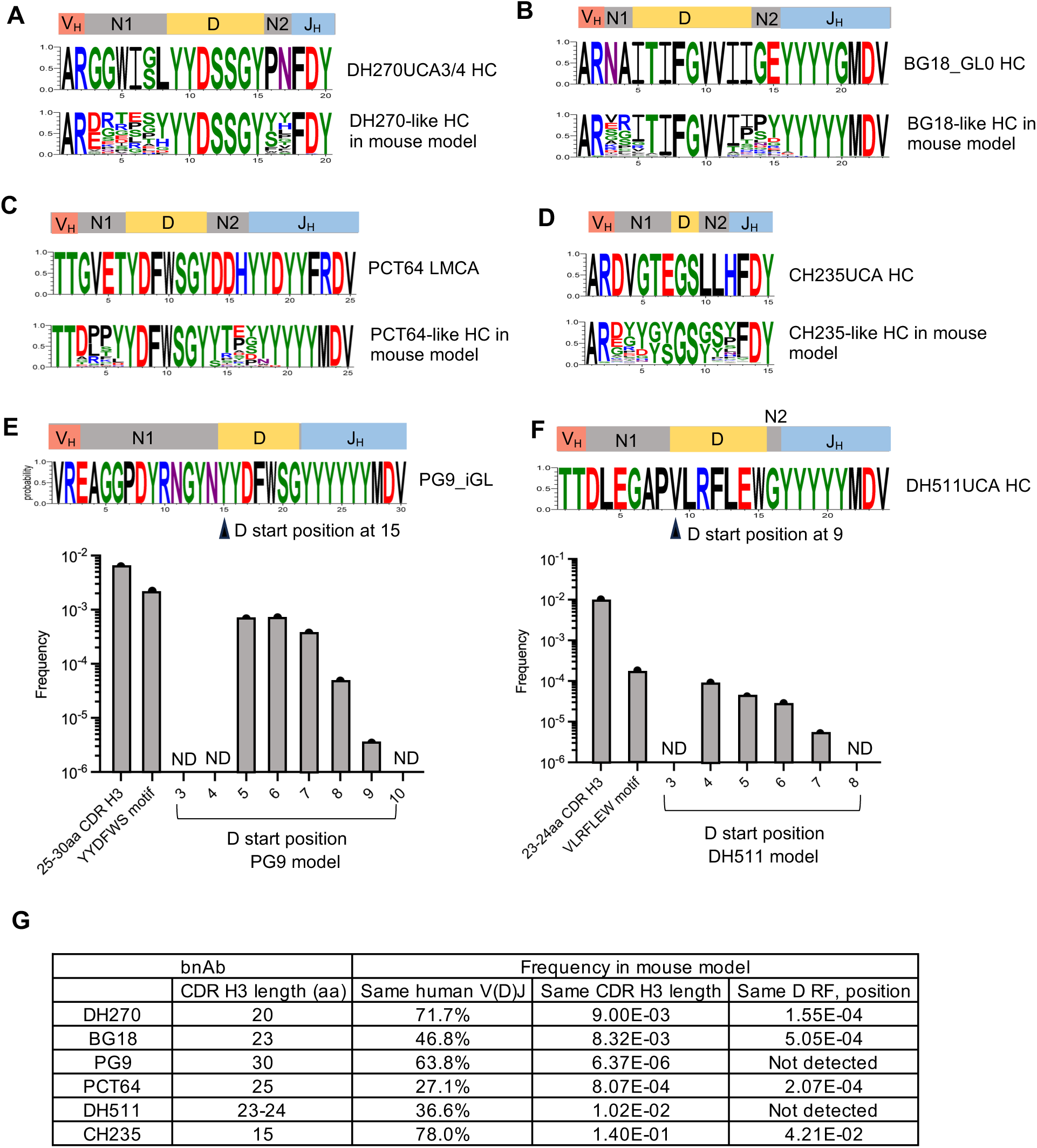
Identification of bnAb-like CDR H3s in the IgH repertoires of the mouse models. (A-D) bnAb-like CDR H3s in the DH270, BG18, PCT64, CH235 models. DH270UCA3/4, BG18_GL0, PCT64LMCA and CH235UCA are inferred precursors for the respective bnAb lineages. In the logo plots, amino acids are represented by standard single letter code, letter color correlates with side chain chemistry, and letter height represents its frequency. The plots were generated with WebLogo 3. (E-F) N1 region analysis of the PG9 and DH511 models. In panel E, “YYDFWS” represents 25-30aa CDR H3s with the D motif. Within this category, D start position is defined as the distance between the 5’ end of CDR H3 and the first “Y” residue in the D motif. The plot in panel F is labeled similarly. Related analysis for 10E8-like CDR H3 is in S7 Fig. (G) Summary of CDR H3 analysis. The frequency of human V/D/J is based on Fig 3F. The frequency for the same CDR H3 length is based on Fig 4A and 4B. The frequency for the same CDR H3 length, same D, reading frame (RF) and position is based on panels A-F of this figure.

We could not identify any PG9-like CDR H3 in the PG9 model. PG9 contains a long 30aa CDR H3 (Fig 5E), which was almost absent in the mouse model (Fig 4B). We relaxed the criteria by widening the CDR H3 range to 25-30aa and found that 0.63% CDR H3s in the PG9 model fell within the 25-30aa range, and 35% of these contained the PG9 D motif (Fig 5E). However, PG9 CDR H3 has an unusually long 12aa N1 region, which positions the 5’ end of D segment at residue 15 of CDR H3 (Fig 5E). In the PG9 mouse model, the 5’ end of the D segment ranged from residues 5-9 of CDR H3s (Fig 5E).

We also did not find any DH511-like CDR H3 in the DH511 model. DH511 bnAbs have 23-24aa CDR H3s. The length was not an issue, since 1% human CDR H3s in the DH511 model were within the range of 23-24aa (Fig 4F). However, like PG9, DH511 CDR H3 also contains a long N1 region. The D motif starts at residue 8-9 of the DH511 CDR H3, but the same motif starts at residues 4-7 of the CDR H3s in the DH511 model (Fig 5F). By contrast, it was easier to identify similar CDR H3s for 10E8, another MPER bnAb that also utilizes hV_H_3-15 and hD3-3 (18). Although CDR H3 lengths of 10E8 (22aa) and DH511 (23-24aa) are similar, the 10E8 CDR H3 has a short 3aa N1 region (S7 Fig). The 10E8-like CDR H3 was found at a frequency of 2.6x10^-4^ (1 in 3846 B cells) in the DH511 model (S7 Fig). The 10E8-like HCs in the DH511 mouse model contained hJ_H_6, whereas 10E8 uses hJ_H_1; but the mismatch should not matter, as the 10E8 immunogen can engage precursors with hJ_H_6 (30, 55).

Since bnAb-like HCs in the mouse models had diverse N regions, their antigen specificities should be heterogeneous. Only a small fraction of the bnAb-like HCs could potentially serve as precursors for a bnAb lineage, especially when CDR H3 constitutes the principal epitope interface. The actual precursor frequency would depend on the immunogen. For example, two versions of DH270 immunogens have been developed. The latest version, CH848 10.17GS, can tolerate more variations in CDR H3 than the prototype, CH848 10.17DT (29, 56). Correspondingly, precursor frequency would likely be higher for CH848 10.17GS than CH848 10.17DT. LC is another variable in precursor frequency but should not be limiting in these mouse models, as shown below.

### IgL repertoire analysis of the mouse models

We employed both flow cytometry and HTGTS-rep-seq to analyze the IgL repertoire of the mouse models. In DH270, BG18 and PG9 models, mCκ has been replaced with hCλ3, which is expressed in association with hVλJλ variable regions. The frequencies of hIgλ_+_ B cells in these mouse models were: 94.9% in the DH270 model, 45.8% in the BG18 model, 97.5% in the PG9 model; the other B cells expressed mIgλ, and there were no mIgκ_+_ B cells, due to substitution of mCκ by hCλ (Fig 6A, 6C, S8A, S8C and S8D Fig). The recombination cassettes in the three mouse models were configured in the same manner but utilized different hVλ segments. Thus, the lower hIgλ_+_ B cell frequency in the BG18 model could be attributable to the hVλ3-25 segment, potentially due to its autoreactivity or poor pairing with HCs. This issue may be related to the overall B cell reduction in the BG18 mouse model (Fig 2). The other abnormality of the BG18 model was increased B1 B cells in spleen (S2D-S2F Fig). The phenotype may not involve hVλ3-25 directly, as the B220^lo^ B1 B cells expressed primarily mIgλ (S8B Fig). Since DH511 LC is hIgκ isotype, it was expressed in conjunction with mCκ and accounted for 87.4% splenic B cells in the DH511 model (Fig 6B, 6C and S8D Fig).

**Fig 6.**
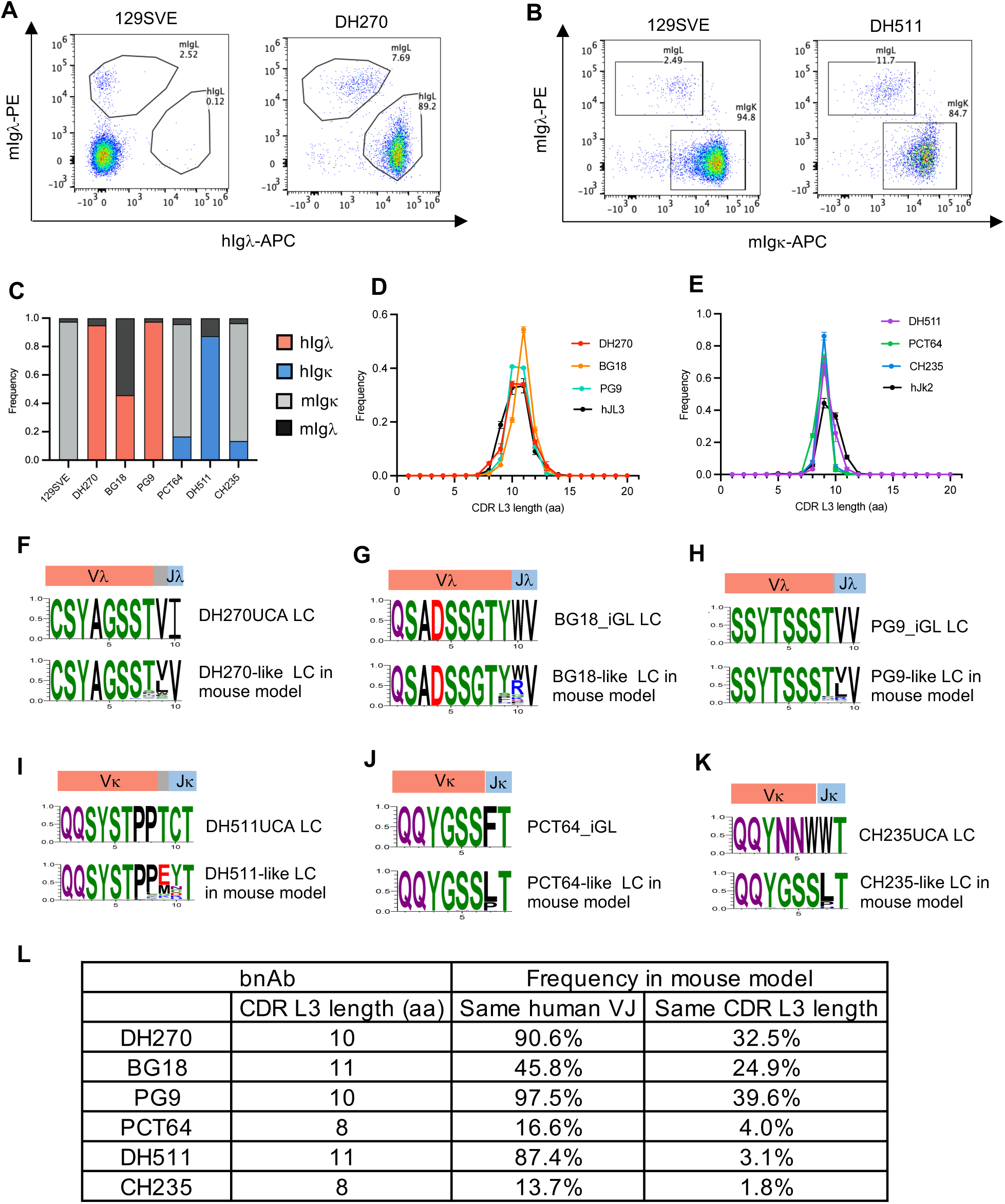
IgL repertoire analysis of splenic B cells in the mouse models. (A-B) Flow cytometric analysis of hIgλ, mIgκ and mIgλ expression in splenic B cells. FACS plots for 129SVE mouse, the DH270 model and the DH511 model are shown here. FACS plots for the other mouse models are in S8 Fig. Gating scheme: lymphocyte>live cell>single cell>B220^hi^>hIgλ/mIgλ or mIgκ/mIgλ. (C) Frequencies of hIgλ_+_, mIgκ_+_ and mIgλ_+_ B cells. The bar graph is based on FACS analysis, as shown in panel A and B, S8 Fig and repertoire analysis in S9 Fig. (D-E) CDR L3 length distribution of hVλ/Jλ and hVκ/Jκ productive rearrangements. The plot for each mouse model is based on the analysis of 3 mice, and error bar represents standard deviation. The data for human is the average of 5 donors, and error bar represents standard deviation. (F-K) bnAb-like CDR L3s in the mouse models. (L) Summary of CDR L3 analysis. The frequency of human V/J is based on panel C. The frequency for the same CDR L3 length is based on panels D-K.

HTGTS-rep-seq analysis confirmed that the humanized Igκ alleles in DH270, BG18, PG9 and DH511 models expressed human variable regions exclusively (S9A-D Fig), consistent with the design of the recombination cassette (Fig 1B). Thus, the frequencies of B cells expressing human variable regions should be equivalent to hIgλ_+_ B cells in DH270, BG18 and PG9 models and mIgκ_+_ B cells in the DH511 model (Fig 6C). As the Igκ alleles in the PCT64 and CH235 models were not based on a recombination cassette, hVκ3-20 accounted for only a fraction of total Igκ rearrangements: 17.4% in the PCT64 model; 13.9% in the CH235 model (S9E-F Fig). Taking mIgλ_+_ B cells into account (S8D Fig), the frequencies of hVκ3-20^+^ B cells in total IgL repertoires were: 16.6% in the PCT64 model; 13.4% in the CH235 model (Fig 6C).

The hVλJλ and hVκJκ rearrangements in the mouse models had similar CDR L3 length distributions as those in human B cells (Fig 6D and 6E). Since IgL rearrangements do not employ D segments, CDR L3 has much lower diversity than CDR H3, as reflected in part by the narrow CDR L3 length distribution (Fig 6D and 6E). As the DH270, BG18 and PG9 antibodies have mainstream CDR L3 lengths for human Igλ LCs: 10-11aa, a substantial fraction of LCs (25-40%) in these mouse model was “bnAb-like”, which was defined as sharing the following properties with bnAb: 1) same hV_L_ and hJ_L_ segment; 2) same CDR L3 length (Figure 6F-6H). The frequency of DH511-like LC was lower (3.1%), because its 11aa length is relatively long for Igκ LCs (Fig 6I). In the PCT64 model, 3.9% LCs qualified as PCT64-like LCs (Fig 6J). As mentioned in the Overview section above, CH235 light chain is composed of hVκ3-15, but the mouse model contained hVκ3-20. Although 1.8% of LCs in the CH235 model contained hVκ3-20 and had 8aa CDR L3, the sequence of these CDR L3s differed substantially from that in CH235UCA, due to divergence of hVκ3-20 and hVκ3-15 at the 3’ end (Fig 6K). Nonetheless, overall homology between hVκ3-20 and hVκ3-15 is 93.4%. hVκ3-20 is used for the 561 class of hV_H_1-46-based CD4 mimic antibodies and contributes to the CDR L3 signature in the paratope (57). Thus, the CH235 model could be useful more broadly for eliciting hV_H_1-46-based CD4 mimic antibodies.

The LC requirements for the bnAbs in these mouse models are not well defined but should be much less demanding than for HCs. In previous immunization experiments, prime immunogens for BG18, PCT64, PG9, CH235 and 10E8 could activate B cells that express bnAb precursor HCs in association with diverse endogenous mouse or rhesus LCs (31, 58–61). However, it is conceivable that further maturation of the bnAb precursors may have more specific LC requirements, and the original human LC of the bnAb would be optimal in later stages of immunization. The V_L_ segment encodes CDR L1, CDR L2 and most of CDR L3. Given the predominant usage of human V_L_ segments in these mouse models, precursor light chains should not be limiting for bnAb induction.

## Discussion

In this work, we generated and characterized six rearranging mouse models that were designed to express rare and variable human bnAb precursors in complex antibody repertoires. Corresponding immunogens have been developed for DH270, BG18, PCT64, PG9 and CH235 bnAbs (21, 23, 29, 31, 56, 58). In the past, we and others have made mouse models to test the immunogenicity of these immunogens (29, 59, 60, 62). In those mouse models, pre-rearranged V(D)J exons for bnAb precursor HC and LC were integrated into the mouse J_H_ and Jκ locus respectively. Because the CDR3s of the pre-rearranged V(D)J exons were fixed, each mouse model expressed a specific bnAb precursor. Immunogen affinity for the precursor can be quantified *in vitro*; precursor frequency is known and can be titrated to desired levels through adoptive transfer (63, 64). Thus, this type of mouse model can provide well controlled environment to address specific questions, such as the affinity threshold for priming a precursor at a given frequency. In contrast, precursors are heterogenous in the rearranging mouse models, and there may also be substantial animal-to-animal variation in precursor pool. Because immunogens are often engineered to engage inferred bnAb precursors, they may not bind as well to divergent precursors in the rearranging mouse models, and their affinities for the heterogenous precursors will also be variable. Since vaccines will face similar difficulties in humans, the rearranging mouse models can be used to assess an immunogen’s performance under such challenging conditions.

We previously made another kind of rearranging mouse model for testing a DH270 immunogen. In that model, the hV_H_1-2 segment and a joined DJ_H_ region of DH270UCA.3 were integrated into the mouse IgH locus (56). During V(D)J recombination, accurate joining of the hV_H_1-2 segment to the DJ_H_ region reconstitutes the original DH270UCA3, whereas imprecise joining creates diversity at the junction to produce a family of DH270UCA3-related HCs. Because the joined DJ_H_ region already includes the N1, D and N2 regions of DH270UCA3 CDR H3, junction diversification is confined to the 5’ end of N1 region, while the present DH270 rearranging mouse model has far more diverse precursors. The hV_H_1-2/joined DJ_H_ model was used to test DH270 immunogen, CH848 10.17GS (56). In that experiment, CH848 10.17GS, was able to activate B cells expressing DH270UCA3 related precursors and promote the acquisition of key mutations of the DH270 lineage. Compared to the previous hV_H_1-2/joined DJ_H_ mouse model, the present DH270 rearranging mouse model should be more stringent for evaluating DH270 immunogens.

The IgH alleles of the BG18, PG9, PCT64 and DH511 mouse models were built on our earlier mouse model that contains the hD3-3 and hJ_H_6 segments (30). The prototype hD3-3/J_H_6 model was used to test 10E8 immunogens. The combination of hD3-3 and hJ_H_6 segments can form long CDR H3, which is needed to reach the recessed MPER epitope. Furthermore, hD3-3 provides critical contact residues for the MPER epitope. Immunization of the hD3-3/J_H_6 mouse model with 10E8 immunogen induced antibodies that contain 10E8 CDR H3 signatures, including long CDR H3. Because the hD3-3/J_H_6 model lacks hV_H_ segments, the 10E8-like CDR H3s were associated with mV_H_ segments. The humanized IgH allele of the present DH511 model includes hV_H_3-15, which is also part of the 10E8 HC and makes direct contacts with the MPER epitope via its CDR H1 and CDR H2 (18). Additionally, IGCRI has been deleted in the DH511 model but not in the hD3-3/J_H_6 model; owing to this difference, hD3-3 was used more frequently in the DH511 model (70.8%) than in the hD3-3/J_H_6 model (16.3%) (30). Given these improvements, the DH511 model should be better than the hD3-3/J_H_6 model for testing the 10E8 immunogen. Although both DH511 and 10E8 utilize hD3-3, these antibodies employ different D motifs to interact with the MPER epitope, and DH511 requires a longer N1 region to position the D motif for epitope contact. In the DH511 model, the closest match for DH511 CDR H3 shifted the D motif toward the 5’end by 2aa (Fig 5F). It is unknown whether such antibody can serve as precursor for DH511 development.

Due to limitation in mouse N region length, PG9-like HC was not found in the PG9 model. PG9-like HCs are also rare in humans. Based on repertoire analysis of 14 human donors, PG9-like HCs were found in only 9 donors, at a frequency of 0.23 per million BCRs (21). As the PG9-like HCs include heterogeneous N regions, the frequency of actual PG9 precursor should be even lower. The ApexGT6 immunogen was designed to target both PG9 and PCT64 precursors. Immunization of rhesus macaques with ApexGT6 elicited diverse V2 apex-specific antibodies (31). These antibodies shared a D motif that can mediate contacts to the V2 apex epitope, and their CDR H3s ranged from 24-33aa. Other than the shared D segment, there was substantial flexibility in D position, N region sequence and length in these elicited rhesus antibodies. Since ApexGT6 appeared capable of engaging a broad range of precursors, it might find suitable targets in the PG9 model as well. However, to contact the C strand of V2 apex, CDR H3 needs to be long enough to penetrate through the glycan shield. The CDR H3 length of the PG9 model can reach the lower range of V2 apex antibodies (24-25aa), but the shortage of “extra-long (>25aa) CDR H3s” would be a limitation for inducing V2 apex antibodies in this mouse model.

VelocImmune and Kymouse are Ig-humanized mice that have incorporated the entire collection of human V, D and J gene segments (52, 65, 66). Based on published Kymouse repertoire data (51), we compared the expression frequencies of human Ig segments that are common to Kymouse and our six rearranging mouse models (S10A Fig). In Kymouse, these Ig segments account for a fraction of the total human IgH repertoire, which includes 38-45 functional V_H_, 23-25 D and 6 J_H_ segments. Furthermore, usage frequency varies substantially for different segments. Relative to the rearranging mouse models, Kymouse contains much lower frequencies of B cells that express relevant human Ig segments and their combinations for the various bnAb lineages in this study (S10 Fig). At such low frequencies, the chance of finding bnAb precursors within the small B cell compartment of the Kymouse is low. The rearranging mouse models are designed to address this limitation by maximizing the expression of bnAb relevant human Ig segments, thereby greatly increasing the associated CDR3 repertoire. The advantage of this approach is evidenced by direct comparison of the efficacies of eOD-GT8 in VRC01 mouse models and Kymouse. The eOD-GT8 immunogen readily elicited VRC01 class precursors in VRC01 mouse models, in which the hV_H_1-2 segment is expressed by about 40% of B cells (34, 37, 67). By contrast, the frequency of hV_H_1-2 is 0.62% in Kymouse (S10A Fig), and induction of the VRC01 class response was much less efficient, due to very low precursor frequencies (68). The performance of the eOD-GT8 immunogen in the VRC01 mouse models proved to be predictive of its success in human clinical trials (69, 70).

Rhesus macaques are widely used for HIV-1 vaccine studies. Genetically, rhesus is obviously closer to human than mouse. Furthermore, outbred rhesus provides a more complex and realistic immune background than inbred mouse to test immunogens. Nonetheless, rhesus and human do have substantial differences in Ig loci. For example, rhesus does not have orthologue for the hV_H_1-2 segment, which contributes directly to epitope contact for DH270 (5) and VRC01 class antibodies (71, 72). As another example, V2 apex antibodies in rhesus uses predominantly the DH3-15 segment (31, 73–76). The RF2 of the DH3-15 segment contains an EDDY motif that is optimal for interaction with the V2 apex C strand. Humans do not have a DH3-15 orthologue. Instead, human V2 apex antibodies primarily use the hD3-3 segment. Rhesus DH3-41 is the closest orthologue of hD3-3, but it is not utilized for rhesus V2 apex antibodies, probably due to strong preference for DH3-15. In these cases, mouse models may be more relevant to human than rhesus in that they contain the exact human Ig segment of interest.

In conclusion, a variety of animal models are now available for preclinical development of HIV-1 vaccines. These animal models differ in bnAb precursor frequencies and total repertoire diversity. Each of these animal models has advantages and limitations. Optimal choice depends on the specific question addressed by a study. With respect to the mouse models described in this study, the main advantage is that each model expresses abundant and diverse human antibodies relevant to a bnAb lineage. The strategy improves the chance of producing rare bnAb precursors within the confines of the small B cell compartment in a mouse. On the other hand, the system lacks the diverse irrelevant human antibodies that may divert immune response off-track during vaccination. Therefore, the model cannot fully assess the immune-focusing ability of an immunogen. Another limitation of the mouse models is the dearth of extra-long CDR H3s for certain types of bnAbs, especially those targeting the V2 apex. In addition to these general issues, the BG18 model exhibited substantial reduction in total B cells and elevation of B1 B cells. It is unknown how the abnormality may impact immunization. Finally, the study describes the baseline unimmunized state of the mouse models. The immune response of these mouse models remains to be evaluated experimentally, and the information presented herein is intended to facilitate their application in immunization studies. In this regard, we have already provided the rearranging mouse models described in this study to several investigators who have relevant immunogens, and we will continue to make the resources available to the HIV-1 vaccine community (see the section of mouse models in Materials and Methods).

## Materials and Methods

### Generation of mouse models

Each mouse model involved several genetic modifications: integration of human Ig segments into the mouse IgH or Igκ loci, IGCRI deletion, insertion of hTdT transgene into the *Rosa* locus. These genetic modifications were introduced into mouse embryonic stem (ES) cells. Site specific integration was achieved by homologous recombination. The cell line was derived from a F1 progeny of 129/Sv and C57BL/6 mouse strains. The IgH alleles of the 129/Sv and C5BL/6 mice differ in sequence. Because homologous recombination has stringent requirements for sequence identity, the difference allowed us to integrate three modifications (hV_H_, hD/J_H_ and IGCRI deletion) into the same IgH allele. The hV_H_, hD-J_H_ and IGCRI deletion are on the 129/Sv allele in the DH270 and CH235 mouse models and on the C57BL/6 allele in the BG18, PG9, DH511, and PCT64 mouse models. The IgH allele choice was based on availability of prior modifications, not any functional considerations. The *Rosa* locus and Igκ locus are on the same chromosome. The Igκ and *Rosa* loci do not have sequence polymorphism between 129/Sv and C57BL/6 alleles to allow allele-specific integration. We generated mouse lines from multiple ES clones and chose the ones where hTdT and the LC recombination cassette co-segregated through breeding. In such mouse lines, the hTdT transgene and LC recombination cassette should reside on the same allele. The co-integration simplifies mouse breeding to reconstitute the complete mouse model.

Each homologous integration utilized a targeting construct. The construct contained the cargo sequence, such as human Ig segment, that was to be integrated into the genome. The cargo sequence was flanked by arms that were homologous to the genomic integration site. The construct also included neomycin resistance gene for positive selection of stable integration and Diphtheria toxin gene for negative selection against random integration. The targeting construct was transfected into ES cell, and stable clones were selected with G418. Genomic DNA was prepared from the stable clones, digested with restriction enzymes that could identify correct integration into correct target site, and screened by Southern blotting. After the correct ES clones were identified, the neomycin resistance gene was deleted via flanking loxP sites by transient expression of cre recombinase in the ES clones; the step eliminated potential inference of the neomycin resistance gene on local transcriptional activity. ES clones in which the neomycin resistance gene was deleted were again identified by Southern blotting. The ES clones were further examined for karyotype. ES clones with normal karyotype (>80% 40 chromosomes) were used for blastocyst injection. S1 Table lists the human Ig segments and associated sequences that have been integrated into the humanized IgH and Igκ alleles. 100bp of local genomic sequences that flank the cago sequences are shown to aid the localization of the integration site.

For each mouse model, we generated one HC ES clone with all the IgH modifications and one LC ES clone with LC recombination cassette plus hTdT transgene. The HC and LC ES clones were injected separately into Rag2 deficient blastocysts. The use of Rag2 deficient blastocysts was to allow lymphocyte analysis at the chimeric mouse stage (77). Since Rag2 is essential for V(D)J recombination, the blastocysts cannot contribute to B and T cells; as a result, all the lymphocytes in the chimeric mice are derived from the injected Rag2 sufficient ES cells. We analyzed B cell phenotypes and antibody repertoires in the chimeric mice. The preliminary analysis was to assess the rearrangement levels of the human Ig segments and their impact on B cell development. After the validation, chimeric mice were bred with C57BL/6 mice for germline transmission. Germline transmission was verified by PCR-based genotyping with tail DNA. For each mouse model, we generated one HC mouse line and one LC mouse line. Subsequently, cross of the HC and LC mouse lines reconstituted the HC/LC mouse model, and both the HC and LC alleles were homozygous in the final model. Homozygous mouse models are maintained in specific pathogen free mouse facility of Boston Children’s Hospital. For distribution of the mouse models, we provide breeding pairs of homozygous mice to recipients and will continue to do so in the near future. We are also in the process of storing the mouse lines through the cryopreservation service of the Jackson Laboratory, which can serve as the long-term repository of the mouse lines. As the mouse lines become widely distributed, interested users may also obtain the mouse lines from investigators who already have them.

### Flow cytometric analysis of splenic B cells

The 129SVE mice were obtained from Taconic (129S6/SvEvTac). All the mice of the mouse models were of mixed 129/Sv and C57BL/6 background and were homozygous for both humanized IgH and Igκ alleles. For analysis of B cell surface markers, single-cell suspension was prepared from spleen. Splenocytes were stained with fluorophore-conjugated antibodies (S2 Table). The stained samples were analyzed on an Attune NxT flow cytometer. FACS plots were generated with FlowJo 10.9.0. The plots are in pseudo-color, and the axis of the plots are biexponential. The plots and gating schemes are described below. Plots are labeled as x/y, and gating for the next plot is in parenthesis. All FACS analysis involve the following steps: FSC-A/SSC-A (lymphocyte gate) > FSC-A/Sytox blue (live cell gate, sytox blue^-^) > FSC-A/FSC-H (single cell gate, diagonal). All the subsequent plots were derived from the single cell gate. 1) total B cells and T cells: CD3/B220 (B cells, B220^+^CD3^-^; T cells, B220^-^CD3^+^). 2) IgM/IgD profile: FSC-A/B220 (B220^hi^) > IgM/IgD (IgM^+^IgD^hi^ gate contains mature follicular B cells). For B220^lo^ cells in BG18 mouse model: FSC-A/B220 (B220^lo^) > IgM/IgD. 3) Follicular and marginal zone B cells: CD93/B220 (mature B cells, B220^hi^CD93^lo^) > CD21/CD23 (follicular B cells, CD21^lo^CD23^hi^; marginal zone B cells, CD21^hi^CD23^lo^). 4) B1 B cell: B220/CD5 (B1 B cells, B220^lo^CD5^+^); B220/CD43 (B1 B cells, B220^lo^CD43^+^). 5) hIgλ, mIgκ, mIgλ B cells: hIgλ/mIgλ, hIgλ_+_ B cells in DH270, BG18 and PG9 mouse models expressed human variable regions; mIgκ/mIgλ, mIgκ_+_ B cells in the DH511 mouse model express human variable region. In PCT64 and CH235 mouse models, only a fraction of mIgκ_+_ B cells expressed hVκ3-20, and the remainder expressed mouse Vκ.

### IgH and IgL repertoire analysis

All the repertoire analyses were done with HTGTS-rep-seq (50). Analysis of mouse models and C57BL/6 mice were done on genomic DNA from splenocytes. For analysis of human B cells, human buffy coat was purchased from BIOIVT. Peripheral blood mononuclear cells (PBMCs) were purified with Ficoll gradient (Cytiva Ficoll-Paque Plus). Naive B cells were positively selected from PBMC with anti-human IgD microbeads with MACS LS column (Milteny Biotech). Genomic DNA was prepared from B cells. For library construction, the genomic DNA was fragmented by sonication in a Bioruptor. Library synthesis was initiated with a primer downstream of the J segment of interest; the J primers are listed in S3 Table. Phusion polymerase (New England Biolab) was used for PCR amplification throughout the procedure. First strand synthesis was carried out through PCR linear amplification. Because the primer was biotinylated at the 5’ end, the first strand was purified with Streptavidin magnetic beads. Adaptor oligonucleotides were ligated to the 3’ end of the immobilized first strand with T4 DNA ligase. The adaptor enabled subsequent PCR amplification of the first strand of the library. The PCR amplification also appended Illumina sequencing adaptors to the library. The PCR amplification also added barcode to each library.

Sequencing was run on Illumina MiSeq, with MiSeq reagent kit v3 600 cycles. Raw sequencing reads (FASTQ) were demultiplexed, based on barcode for each library. The demultiplexed reads underwent deduplication to combine identical reads into one read. Deduplicated reads were aligned to a test sequence that is internal to the 3’ end of the PCR primer that was used to amplify and append sequencing adaptor to the library. The reads that aligned to the internal sequence were correctly primed on-target reads. The on-target reads were aligned to a test sequence that was immediately upstream of the J segment. The sequence should be eliminated by V(D)J recombination to the J segment. The reads that did not contain the test sequence had undergone V(D)J recombination to the J segment. These rearranged reads were aligned to a database that contained mouse Ig segments and relevant human Ig segments. The alignment was based on IgBlast. The output included the sequence of the V(D)J rearrangement, identity of the V/D/J segment, productive or nonproductive, and CDR sequence and length. The IgH and IgL repertoire analysis plots were based these outputs.

The analysis of human Ig usage in Kymouse was based on published data (51). The data sets were IGM sequences from naive Kymouse 1, 2, 3, 4, 5, 6, and 7. The usage frequencies of the Ig segments and V(D)J rearrangements in S10 Fig are the average value from 7 Kymouse. Only productive rearrangements were used for the analysis.

## Acknowledgments

We thank Thandi Onami at the Gates Foundation for regular discussions, suggestions and administering the project. We also thank Barton Haynes and collaborators at Duke CHAVD for inputs into the work. We thank Kerstin Johnson, Nicole Manfredonia and Lucas Vieira Francisco for assistance with mouse work.

## Author contributions

**Conceptualization:** Ming Tian, Frederick W. Alt

**Formal Analysis:** Ming Tian, Hwei-Ling Cheng, Jillian Davis, Lily M. Thompson

**Funding acquisition:** Frederick W. Alt, Ming Tian

**Investigation:** Ming Tian, Hwei-Ling Cheng, Jillian Davis, Lily M. Thompson, Aimee Chapdelaine Williams, Marie-Elen Tuchel, Audrey Yin

**Methodology:** Ming Tian, Hwei-Ling Cheng, Jillian Davis, Lily M. Thompson

**Project Administration:** Hwei-Ling Cheng, Ming Tian, Frederick W. Alt

**Resources:** Frederick W. Alt

**Software:** Lawrence Jianqiao Hu, Xin Lin, Adam Yongxin Ye

**Supervision:** Ming Tian, Frederick W. Alt

**Validation:** Ming Tian, Hwei-Ling Cheng, Jillian Davis, Lily M. Thompson

**Visualization:** Ming Tian, Lily M. Thompson

**Writing – Original Draft Preparation**: Ming Tian, Frederick W. Alt

**Writing – Review & Editing:** Ming Tian, Jillian Davis, Lily M. Thompson, Frederick W. Alt

## Ethics Statement

All the mouse works were approved by the Institutional Animal Care & Use Committee (IACUC) of Boston Children’s Hospital (protocol #00002005).

## Data availability statement

Library sequencing for HTGTS-rep-seq analysis was performed on Illumina Miseq. The raw sequencing reads, in FASTQ format, were deposited in Sequence Read Archive. The accession numbers for the mouse and human HTGTS-rep-seq data are: PRJNA1439958 and PRJNA1439838, respectively. All other data for this work are included in the manuscript.

## Supporting information

**S1 Fig.**
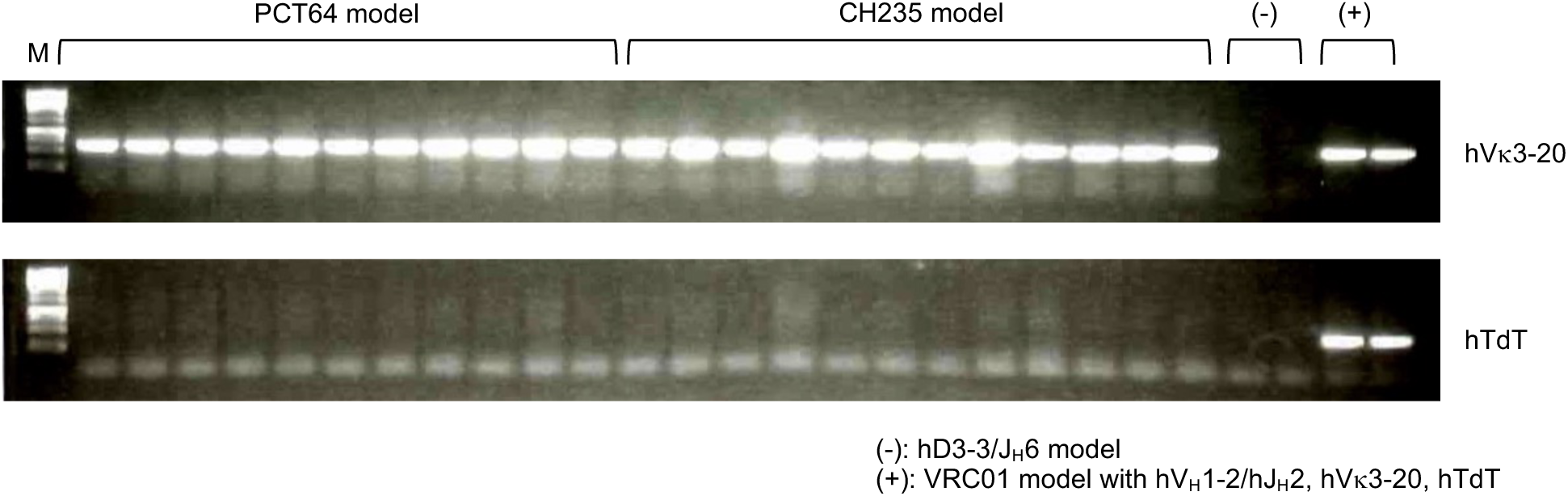
Genotyping of hVκ3-20 and hTdT in the PCT64 and CH235 models. M: 1kb DNA ladder. Each following lane contains the genotyping result of one mouse.

**S2 Fig.**
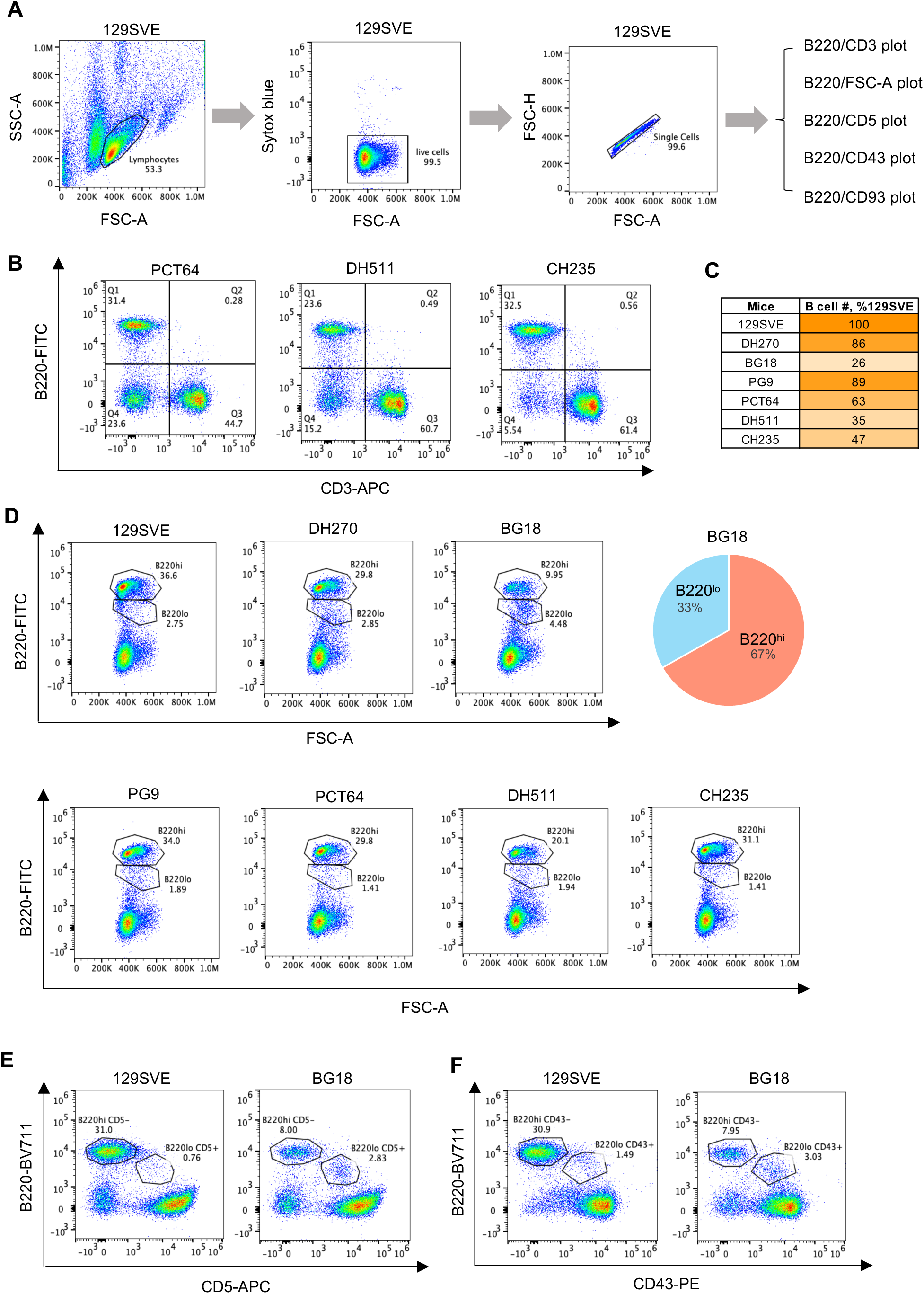
Flow cytometric analysis of B cells and T cells in the spleen of the mouse models. (A) Gating scheme for downstream plots in Fig 2A and panels B-E of this figure. The analysis of 129SVE splenocytes is shown here as an example. Splenocytes from the mouse models were analyzed in the same way. (B) Quantification of total B and T cells. These plots supplement Fig 2A. (C) Comparison of splenic B cell numbers in different mouse models. To calculate B cell numbers, the frequency of B220^+^ B cells was multiplied with total splenocyte number. The average value for each mouse model was divided by that for the 129SVE mouse to yield the values in the table. (D) Separation of B220^hi^ and B220^lo^ populations according to B220 staining and size (FSC-A). The B220^hi^ gate proceeds the IgM/IgD plot (Fig 2B and S3A Fig). The pie chart shows the distribution of B220^hi^ and B220^lo^ populations in the BG18 model. (E-F) Expression of B1 B cell marker (CD5 and CD43) in the B220^lo^ population. B1 B cells are B220^lo^CD5^hi^CD43^hi^.

**S3 Fig.**
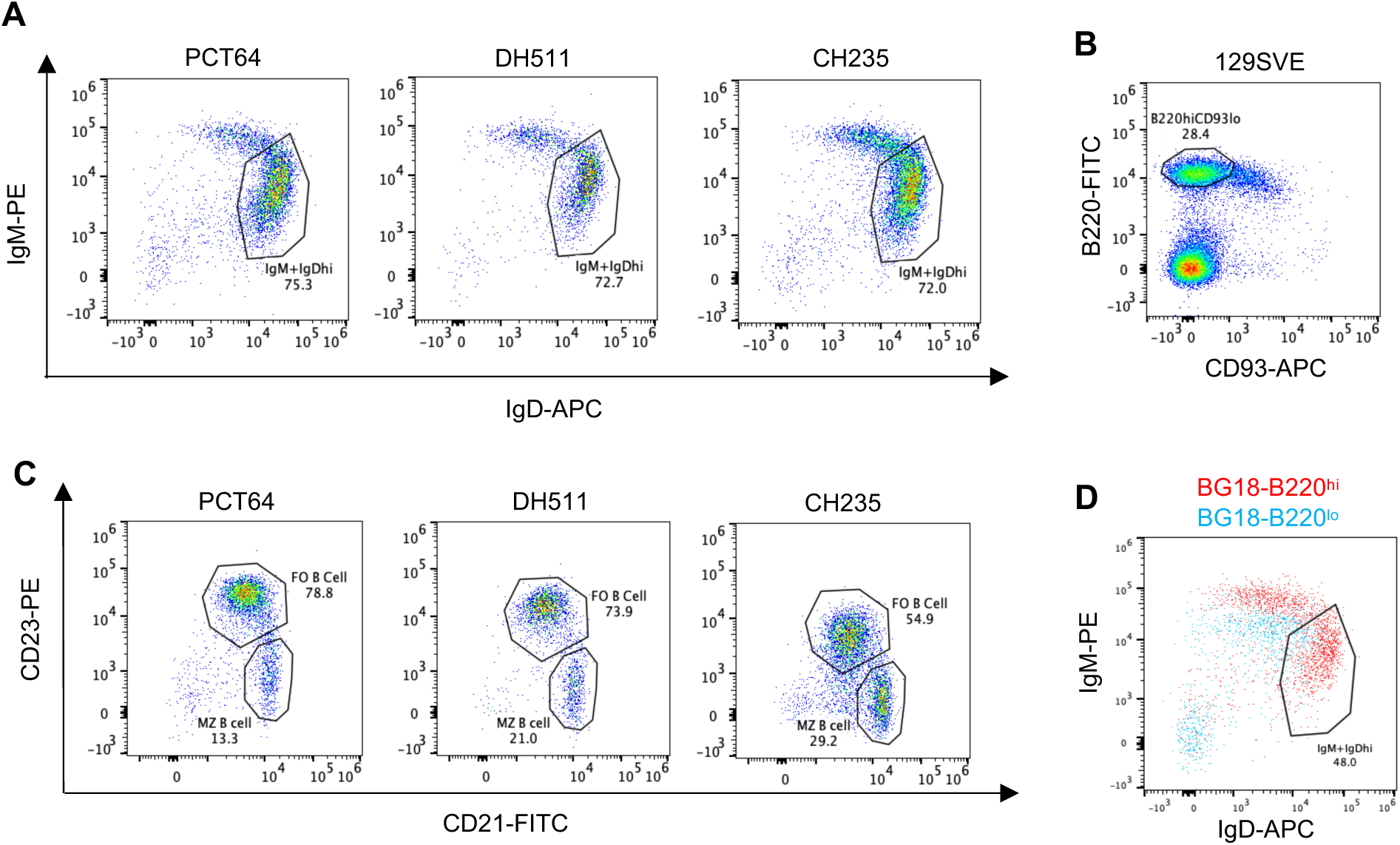
Flow cytometric analysis of IgM/IgD profile, follicular B cells and marginal zone B cells in the spleen of the mouse models. (A) IgM/IgD staining of B220^hi^ cells. These FACS plots supplement Fig 2B. (B) Gating for mature B cells (B220^+^CD93^lo^) in the CD23/CD21 plots in Fig 2C and panel D of this figure. 129SVE splenocytes were analyzed in these FACS plots. The same gating scheme was applied to the analysis of all the mouse models. (C) Follicular and marginal zone B cells. These FACS plots supplement Fig 2C. (D) Overlay of IgM/IgD staining of B220^lo^ and B220^hi^ cells in the BG18 model.

**S4 Fig.**
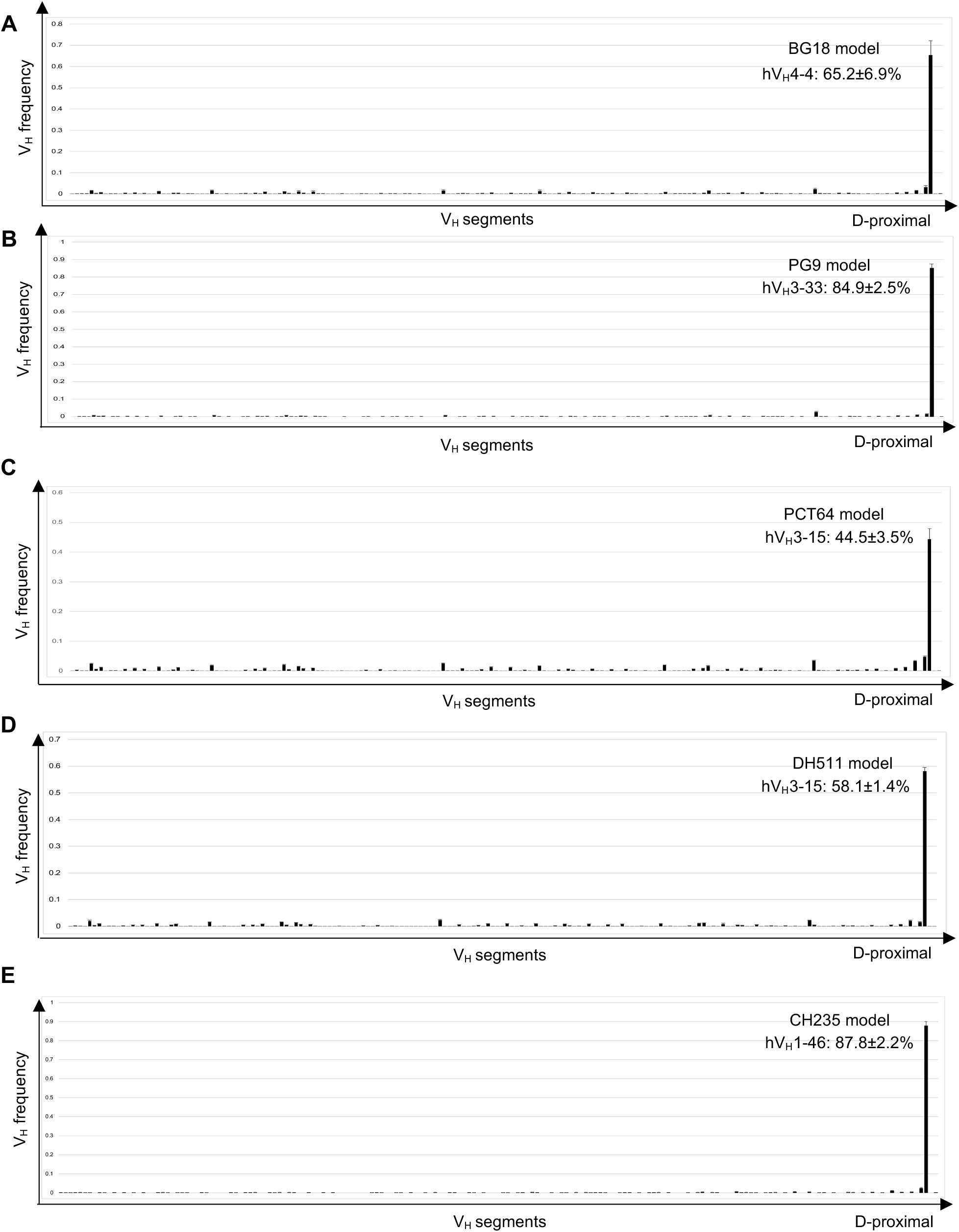
V_H_ usage of productive IgH rearrangements in splenic B cells of the mouse models. (A-E) V_H_ usage frequencies in the BG18, PG9, DH511, PCT64 and CH235 models. These plots supplement Fig 3A. The data are summarized in Fig 3B.

**S5 Fig.**
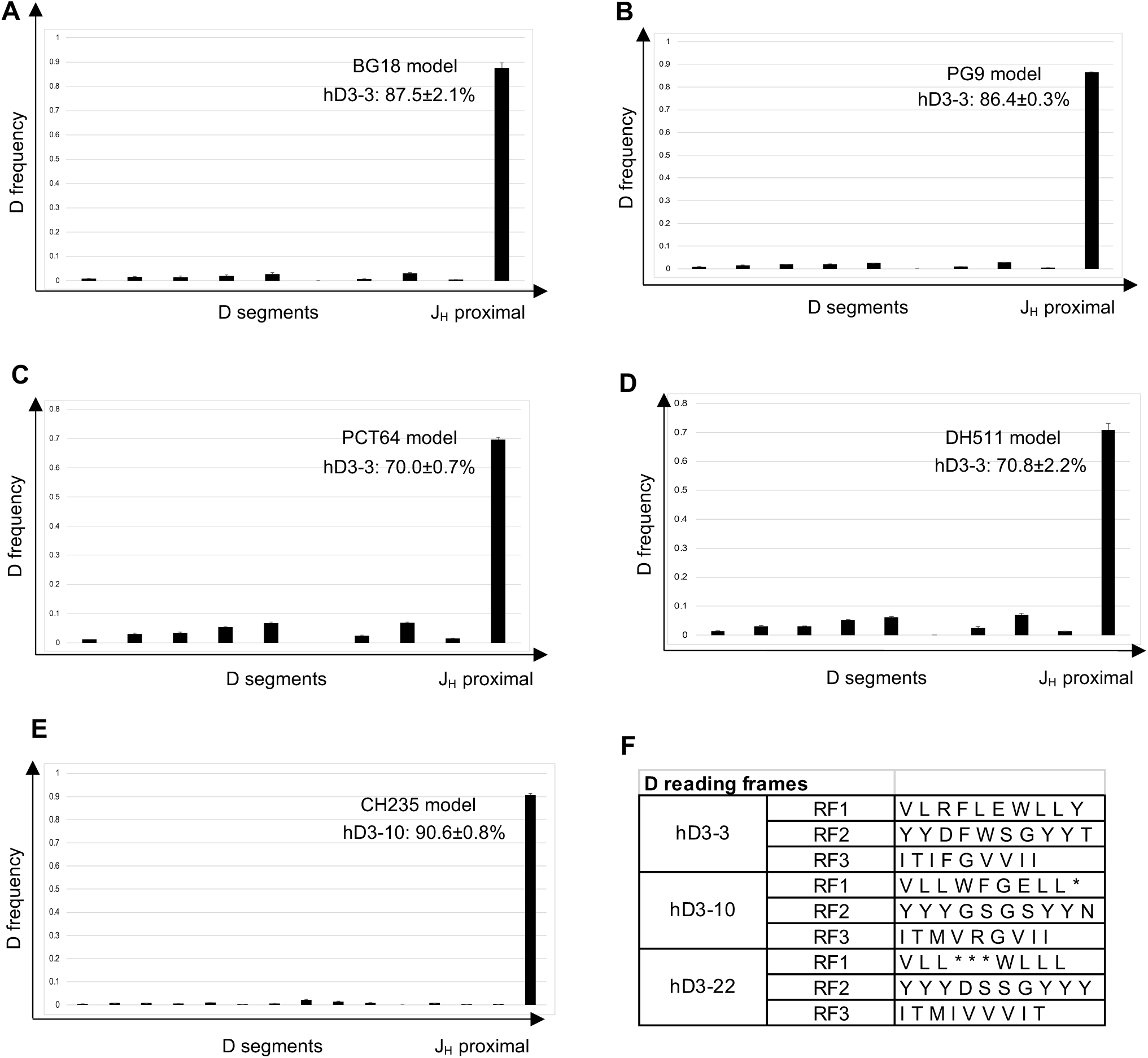
D usage of productive IgH rearrangements in splenic B cells of the mouse models. (A-E) D usage frequencies in the BG18, PG9, DH511, PCT64 and CH235 models. The plots show D usage frequencies in productive rearrangements involving hV_H_ and hJ_H_. These plots supplement Fig 3C. The data are summarized in Fig 3D. (F) D reading frames. The table lists the sequences of each D reading frame and supplements Fig 3E.

**S6 Fig.**
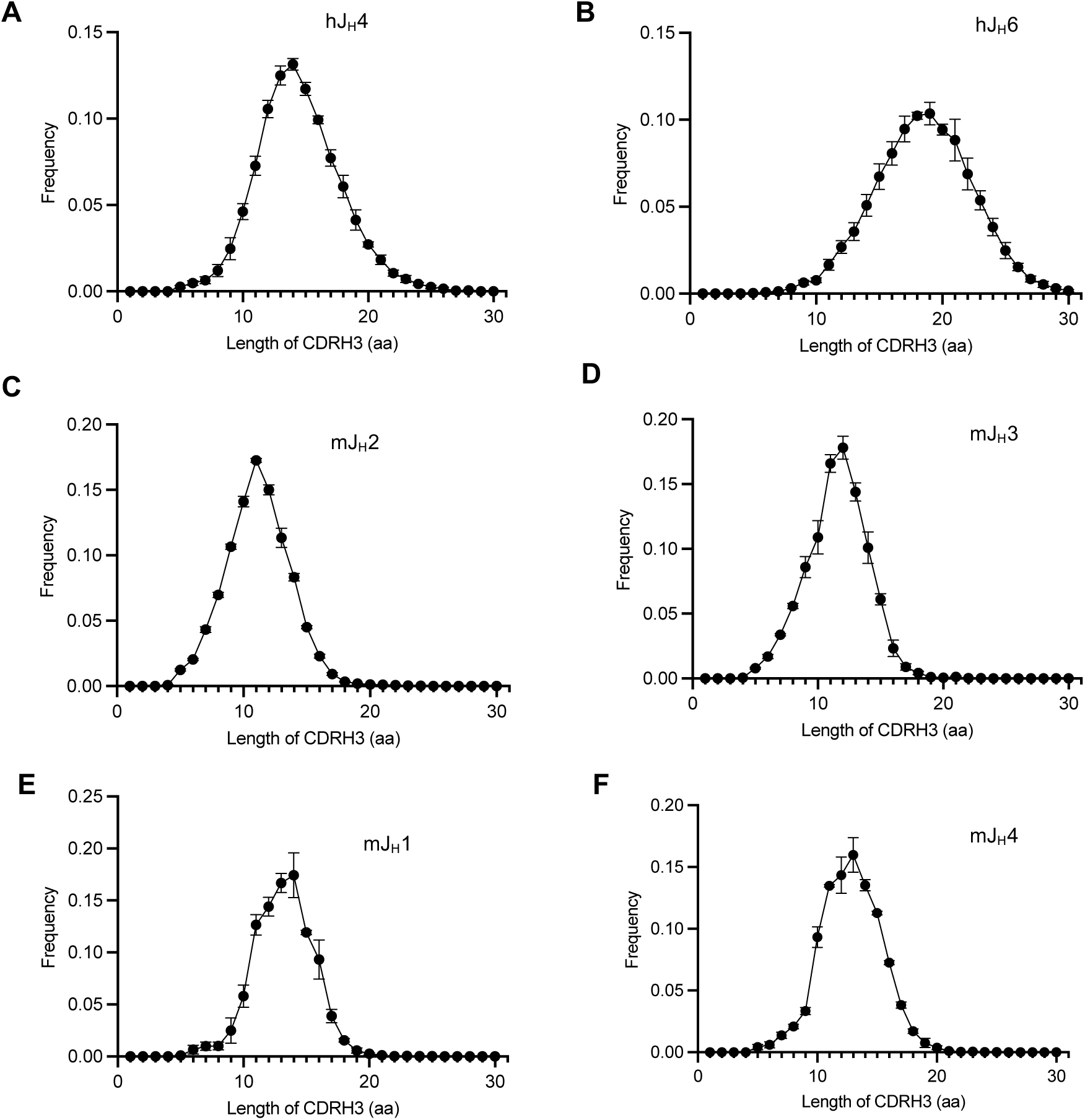
CDR H3 length distribution of human peripheral blood B cells and C57BL/6 mouse splenic B cells. The human data were generated by HTGTS-rep-seq analysis of naive B cells (IgD^+^) from peripheral blood. The plots are average of 5 human donors; error bar represents standard deviation. The mouse data were generated by HTGTS-rep-seq analysis of splenic B cells of C57BL/6.SJL mice. The plots are average of 3 mice; error bar represents standard deviation. (A-B) CDR H3 length of human V_H_/D/J_H_4 and V_H_/D/J_H_6 productive rearrangements. (C-F) CDR H3 length of mouse J_H_1, J_H_2, J_H_3 and J_H_4-based productive rearrangements.

**S7 Fig.**
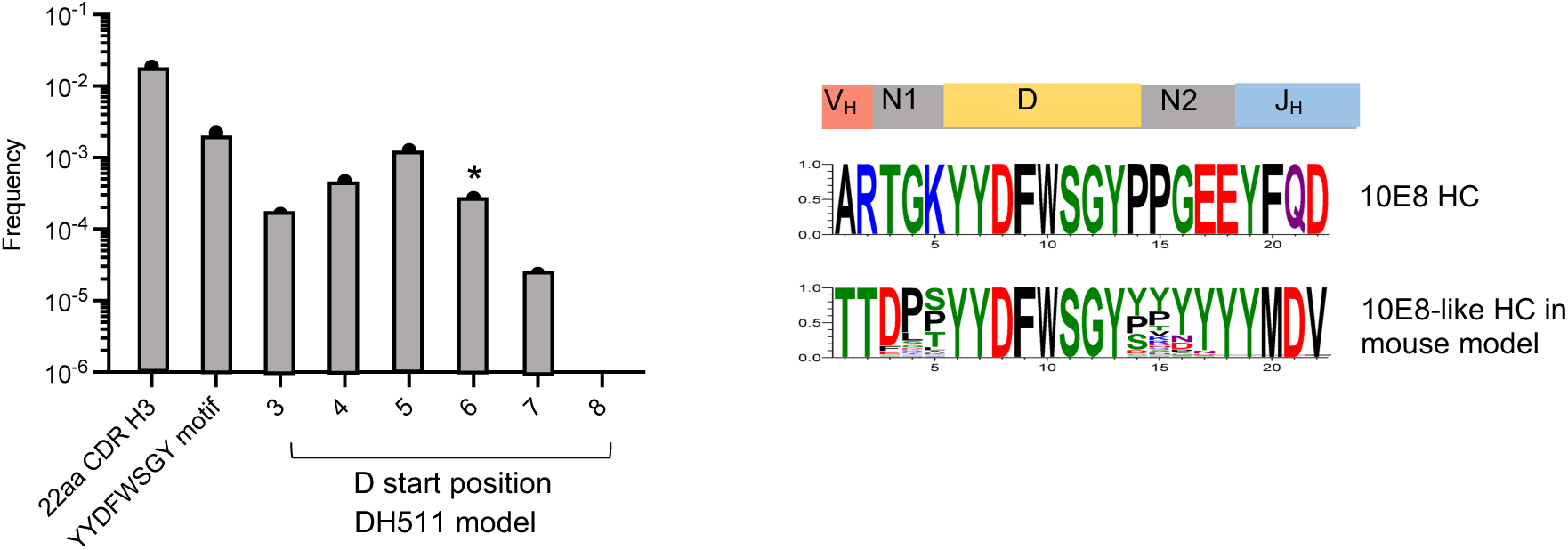
Analysis of 10E8-like CDR H3 in DH511 mouse model. D start position in 10E8 CDR H3 is marked with *.

**S8 Fig.**
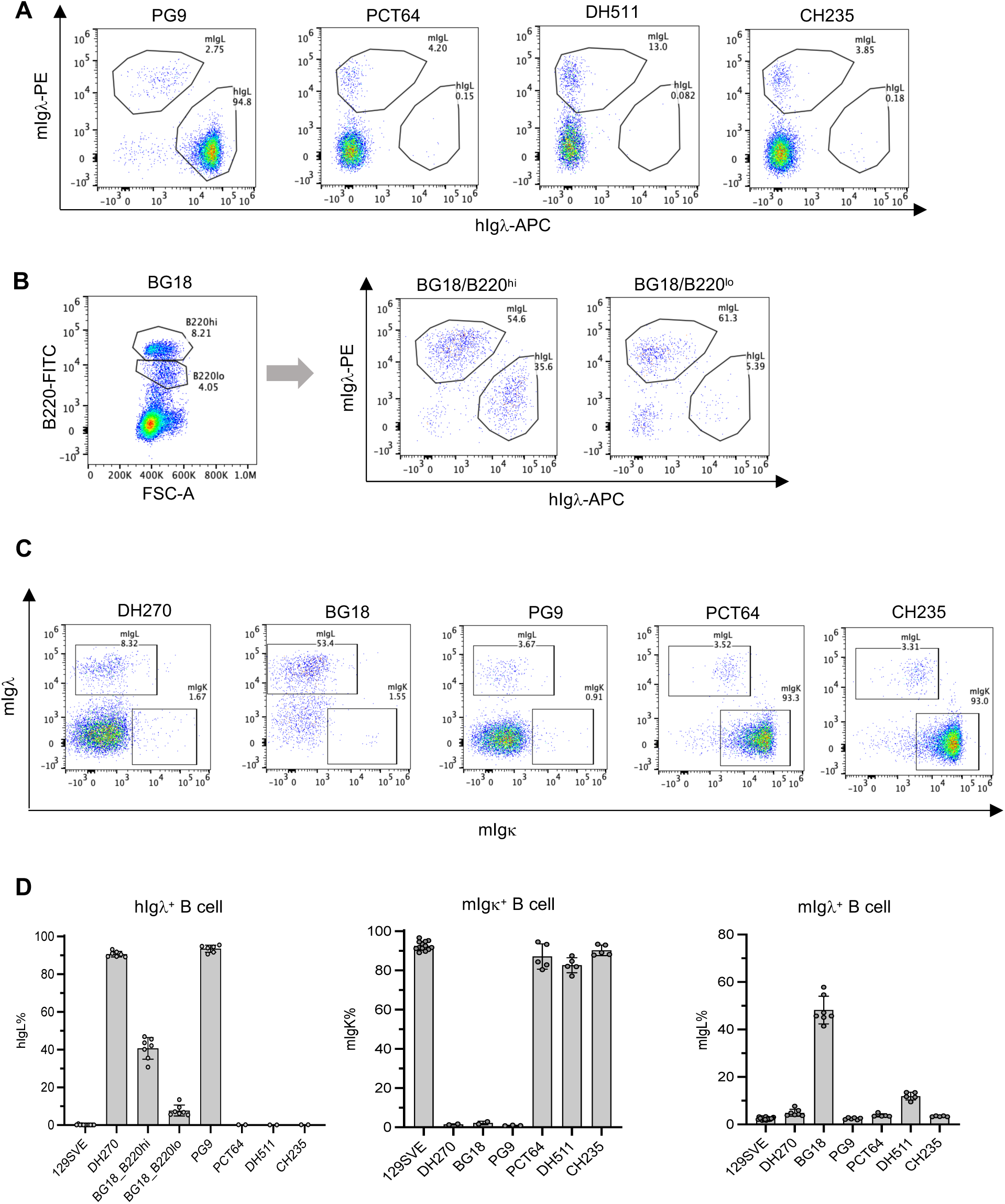
IgL repertoire analysis of mouse models. (A) Flow cytometric analysis of hIgλ and mIgλ expression in splenic B cells. The data supplement Fig 6A. (B) hIgλ/mIgλ expression in B220^hi^ and B220^lo^ population in the BG18 mouse model. (C) Flow cytometric analysis of mIgκ and mIgλ expression in splenic B cells. The data supplement Fig 6B. (D) Frequencies of hIgλ_+_, mIgκ_+_ and mIgλ_+_ B cells. The bar graphs summaries the FACS analysis in Fig 6A, 6B and panels A-C of this figure. Each dot represents one mouse. The height of the bar equals the average, and error bar corresponds to standard deviation.

**S9 Fig.**
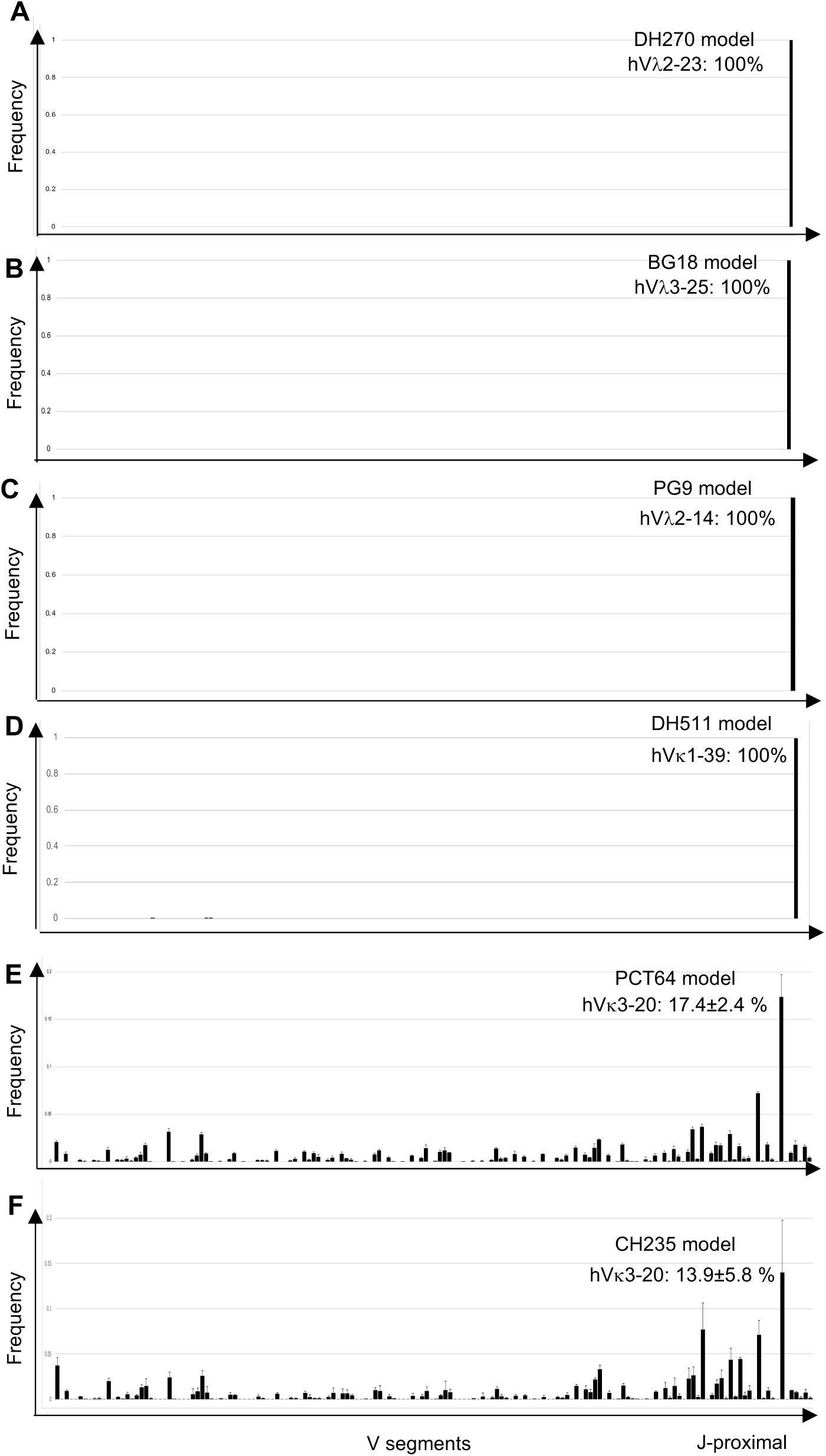
V_L_ usage among productive rearrangements on the humanized Igκ allele. (A-F) The plots show the results of HTGTS-rep-seq analysis on splenic B cells of the mouse models. The plots are based on the average of three mice; error bar represents standard deviation.

**S10 Fig.**
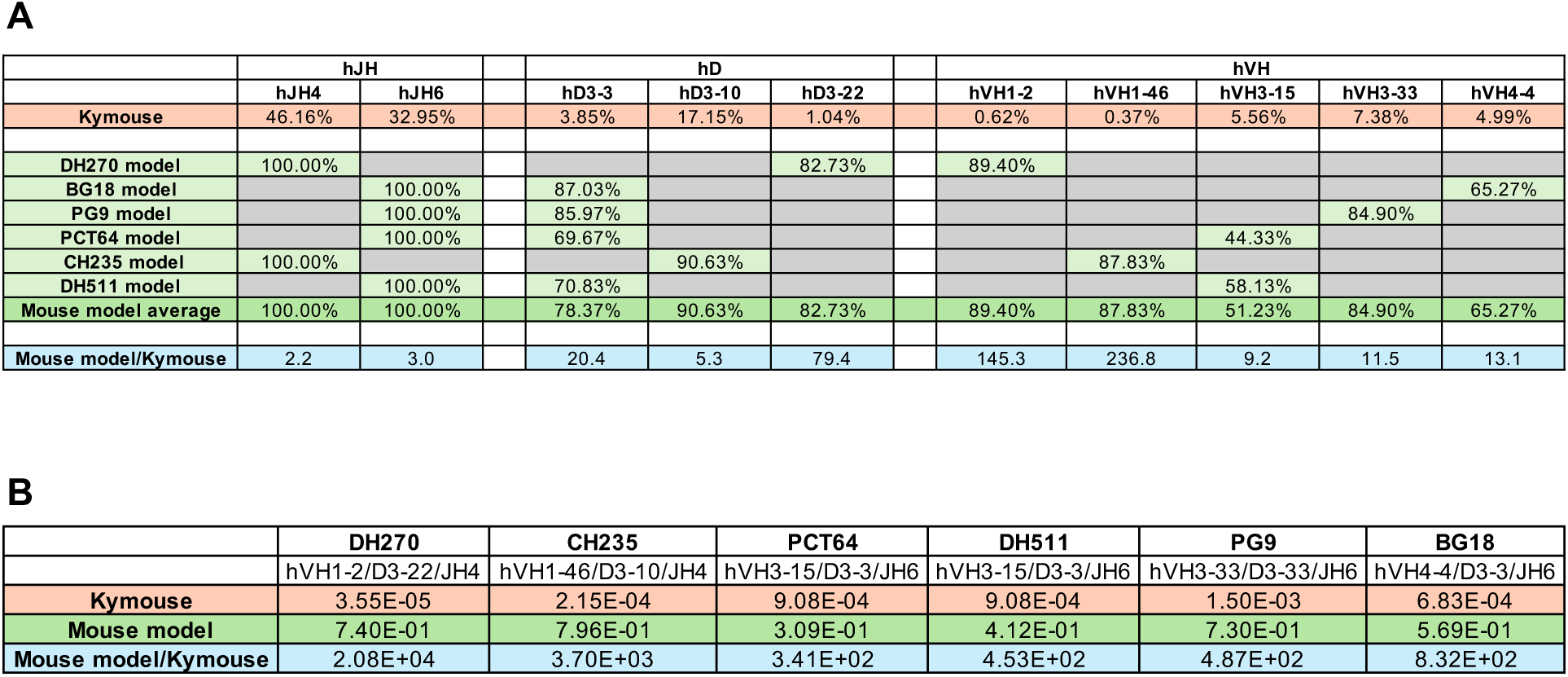
Comparison of human Ig segment usage in Kymouse and rearranging mouse models. (A) The table lists the percentage of productive V(D)J rearrangements that contain the indicated human Ig segments. These Ig segments are utilized in both the rearranging mouse models in this work and in Kymouse. The data for the rearranging mouse models are based on Fig 3B and 3D (the mean value for each mouse model). The values for Kymouse are based on analysis of published data (see Material and Methods). (B) The table lists the frequencies of productive V(D)J rearrangements that have the same Ig segment compositions as the six bnAb lineages. The data for the rearranging mouse models are based on Fig 3F (the mean value for each mouse model). The values for Kymouse are based on analysis of published data (see Material and Methods).

**S1 Table.** The sequences of the human Ig segments and adjacent regions in the mouse models.

**S2 Table.** Antibodies used in flow cytometric analysis.

**S3 Table.** J primers used in HTGTS-rep-seq analysis.

